# Mutations in disordered regions cause disease by creating endocytosis motifs

**DOI:** 10.1101/141622

**Authors:** Katrina Meyer, Bora Uyar, Marieluise Kirchner, Jingyuan Cheng, Altuna Akalin, Matthias Selbach

**Author notes:** Corresponding author: Matthias Selbach Max Delbrück Centrum for Molecular Medicine Robert-Rössle-Str. 10, D-13092 Berlin, Germany.

## Abstract

Mutations in intrinsically disordered regions (IDRs) of proteins can cause a wide spectrum of diseases. Since IDRs lack a fixed three-dimensional structure, the mechanism by which such mutations cause disease is often unknown. Here, we employ a proteomic screen to investigate the impact of mutations in IDRs on protein-protein interactions. We find that mutations in disordered cytosolic regions of three transmembrane proteins (GLUT1, ITPR1 and CACNA1H) lead to an increased binding of clathrins. In all three cases, the mutation creates a dileucine motif known to mediate clathrin-dependent trafficking. Follow-up experiments on GLUT1 (SLC2A1), a glucose transporter involved in GLUT1 deficiency syndrome, revealed that the mutated protein mislocalizes to intracellular compartments. A systematic analysis of other known disease-causing variants revealed a significant and specific overrepresentation of gained dileucine motifs in cytosolic tails of transmembrane proteins. Dileucine motif gains thus appear to be a recurrent cause of disease.

---

Genome sequencing technologies have greatly facilitated the discovery of human protein variants. In many cases it is not known if such variants cause disease, and even when associations have been discovered, determining the mechanisms by which this happens remains a major challenge ^1^. Most disease-causing missense mutations affect evolutionarily conserved amino acids within structured regions of a protein and destabilize its structure ^2,3^. However, over 20% of human disease mutations occur in so called intrinsically disordered regions (IDRs) ^4^. Contrary to the traditional understanding of protein structure and function, it is now clear that IDRs represent a functionally important and abundant part of eukaryotic proteomes ^5,6^. Yet, since IDRs lack a defined tertiary structure and are typically poorly conserved, the classical structure-function paradigm cannot explain how mutations in IDRs cause disease.

One way to approach this issue is by analyzing protein-protein interactions (PPIs), which can help to understand how mutations cause disease ^7,8^. The impact of PPIs on disease is highlighted by the enrichment of missense mutations on interaction interfaces of proteins associated with the corresponding disorders ^9^. Moreover, comparing the interaction partners of wild-type proteins and their disease-associated variants can reveal disease mechanisms ^10,11^. We therefore sought to systematically investigate how mutations in IDRs affect PPIs.

IDRs often harbour short linear motifs (SLiMs) which mediate their function ^12,13^. These SLiMs typically fall into two major classes -- motifs that mediate interactions with globular domains and/or motifs which harbor posttranslational modification sites ^14^. Mutations in IDRs can cause disease by disrupting such motifs or by creating novel ones. A number of examples of such pathogenic changes in motifs have been reported in the literature ^15–18^. Additionally, computational studies have revealed that pathogenic mutations often target SLiMs ^19–21^. Despite these insights, however, there has not yet been a systematic experimental analysis of the way disease-causing mutations in IDRs affect interactions. One reason for this is that the small binding area between SLiMs and cognate domains results in low binding affinities, which makes it difficult to study these interactions ^22^.

## A peptide-based interaction screen on disease-causing mutations

We reasoned that quantitative interaction proteomics with immobilized synthetic peptides should enable us to systematically assess the impact of mutations in IDRs. Such peptide pull-downs can maintain specificity even with low affinity interactions ^23^. We therefore designed a scalable proteomic screen that employs peptides synthesized on cellulose membranes (Fig. 1 A). These membranes carry peptides with a length of 15 amino acids that correspond to IDRs in both the wild-type and mutant form. Membranes are incubated with cell extracts to pull-down interacting proteins. After washing, peptide spots are excised and the proteins associated with them are identified and quantified by shotgun proteomics. The main challenge in such interaction screens is to distinguish specific interaction partners from non-specific contaminants ^24–27^. We addressed this challenge through the use of two levels of quantification. First, two replicates of a pull-down with a specific peptide sequence are compared to all other peptide pull-downs via label-free quantification (LFQ) ^28^. This LFQ-filter selects proteins that bind specifically to a given peptide. Secondly, the screen uses SILAC-based quantification ^29^ to identify differential interaction partners of the wild-type and disease-causing form of a peptide. This requires incubating the two replicates of the membrane with cell lysates that have been differentially SILAC labeled. Wild-type peptide spots from the heavy pull-down are combined with spots from the light pull-down that correspond to the mutant forms of the same peptide, and *vice versa*. SILAC ratios give a measure of the degree to which each particular mutation affects a specific interaction.

**Fig. 1:**
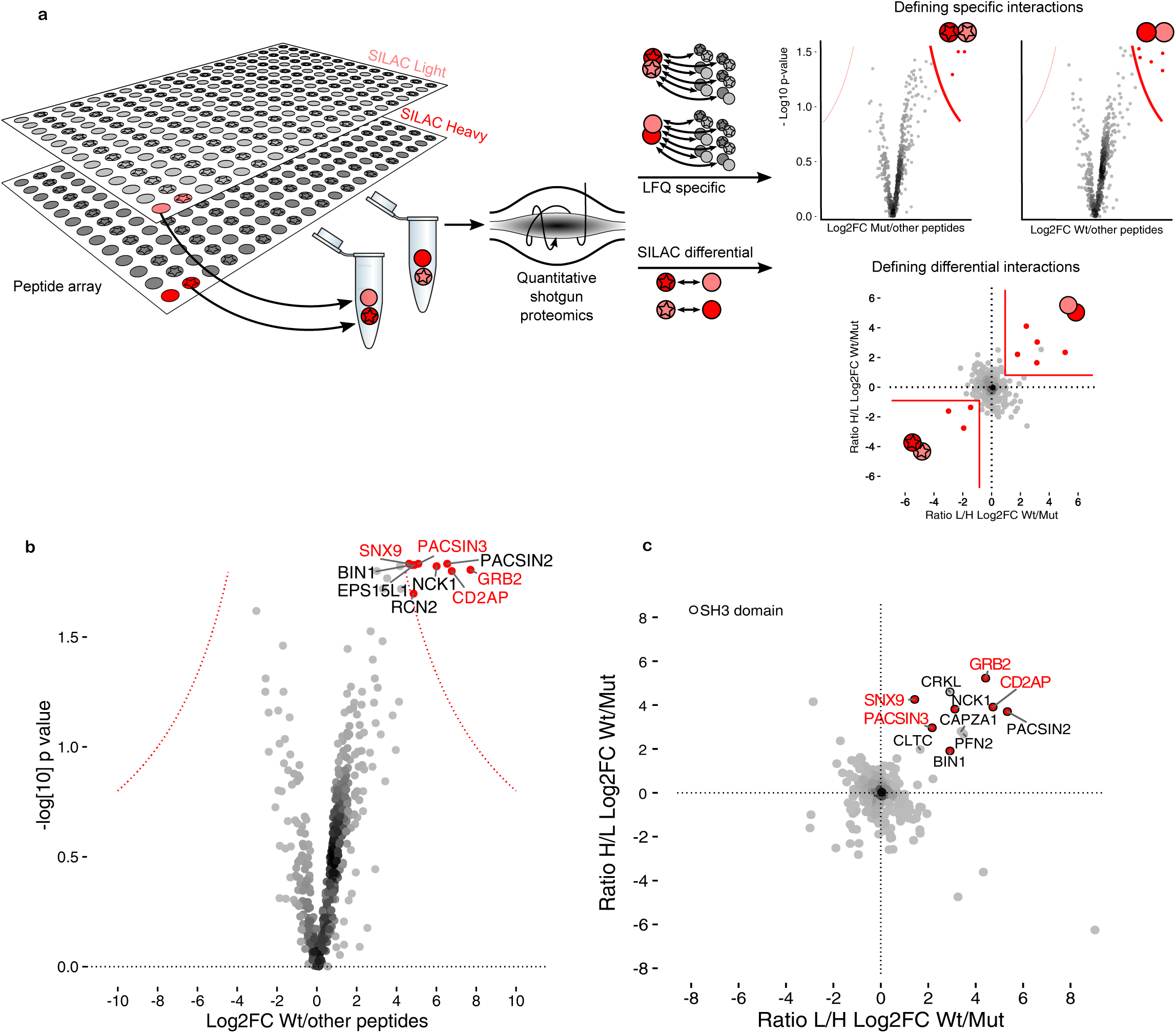
Quantitative interaction screen with disease-associated disordered regions. **A**, Cellulose membranes with synthetic wild-type (circles) and mutated (stars) peptides are incubated with lysate from light (light red) or heavy (signal red) SILAC labeled cells to pull-down interacting proteins. Spots are excised, corresponding wild-type/mutant pairs are combined and analyzed by quantitative shotgun proteomics. Label free quantification (LFQ) identifies specific interactors by comparing both replicates to all other pull-downs. SILAC identifies differential binders by directly comparing corresponding wild-type and mutant pairs. **B**, **C** Results for a SOS1-derived peptide with a proline-rich motif as a benchmark. **B**, Volcano plot from the LFQ data for wild-type SOS1. Specific binders are shown as red dots. 4 out of 5 known binders (red gene names) are detected. **C**, SILAC log2 fold changes for differential binders of the wild-type and mutant SOS1 peptide. Proteins with SH3 domains are shown with black outlines.

For the screen we selected 128 mutations in IDRs which are known to cause neurological diseases (Fig. S1, Table S1). We included a peptide from an IDR in the SOS1 protein that contains a proline-rich motif by which it is known to recruit several specific binders via their SH3 domains ^23^. We analyzed the 2 × 129 pull-down samples by using high-resolution shotgun proteomics in 45 min runs, resulting in a total measurement time of about eight days. Replicates of the same peptide clustered with each other with a median correlation coefficient of 0.87 (Pearson’s R), indicating good reproducibility (Fig. S2). The LFQ data identified nine specific interactors of the SOS1 peptide, including four of the five that were previously known (Fig. 1 B). In the corresponding SILAC data, seven of the nine LFQ-specific binders show preferential binding to the wild-type compared to the mutant which has a disrupted proline-rich motif (Fig. 1 C). Importantly, all interactors that are both specific (LFQ) and differential (SILAC) contain SH3 domains. To further assess the relationship between peptide motifs and cognate domains we also analyzed all pull-downs combined. We found that mutations which disrupt a predicted SLiM in the peptide tend to reduce binding of proteins with cognate domains (Fig. S3). Conversely, the gain of a SLiM in a peptide tends to increase its binding to proteins with matching domains. In summary, these data demonstrate that our screen efficiently detects how mutations in IDRs affect interactions mediated by SLiMs.

## A quantitative interaction network for disease-associated IDRs

Individual pull-downs typically led to the identification of ~ 400 proteins. If all of these proteins were considered specific binders, this would correspond to ~ 400 binary interactions per pulldown and a total of more than 100,000 interactions. However, since many proteins are in fact background binders, we applied our quantitative filters with cut-offs derived from the SOS1 control (Fig. 2 A). About half of the 2x128 peptides showed at least one specific binder according to the LFQ-filter (Fig. S4). Applying the LFQ-filter dramatically reduced the total number of interactions to 618. All of these 618 interactions are specific for the wild-type and/or the mutant form of a peptide as compared to all other peptides in the screen (Table S2). However, not all of these specific interactions are differential, i.e. affected by the mutation. Therefore, we next applied the SILAC-filter, which led to a final list of 180 differential interactions (Table S3). 111 of these interactions are lost through mutations in the peptide, while 69 are gained. Of note, since pull-downs can also capture indirect binders, not all of these interactions are necessarily direct.

**Fig. 2:**
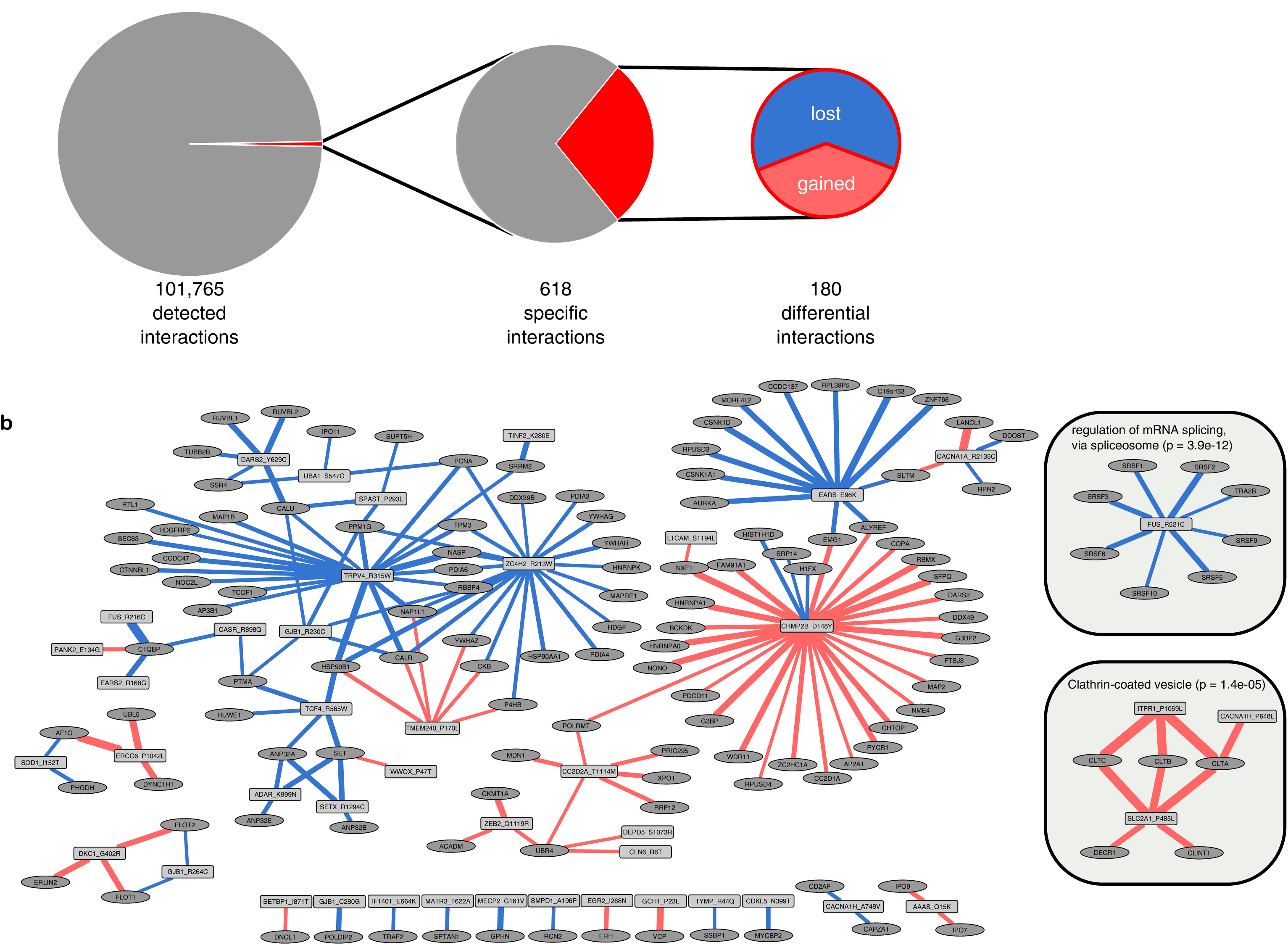
Differential interactors of wild-type and mutant IDRs. **A**, Quantitative filters to select specific and differential interactions. Only a minor fraction of all detected interactions is specific (LFQ filter). Moreover, only a fraction of specific interactions is differential (SILAC filter), i.e. show preferential binding to the wild-type or mutant form of a peptide. Mutation-induced interaction losses are more frequent than mutation-induced gains. **B**, Network of all differential interactions. Peptides (rectangles) and interacting proteins (ovals) are presented as nodes. The edges indicate preferential binding to the wild-type (blue) or mutant (red) form of a peptide (edge width indicates SILAC ratios). Highlighted subnetworks are enriched in splicing regulators and clathrin-coated vesicle proteins (see text).

To provide an overview of the data we displayed all of the differential interactions as a network (Fig. 2 B). This revealed that several wild-type or mutant peptides shared differential interactors, suggesting functional similarities. Moreover, subnetworks were enriched in specific gene ontology terms (Fig. S5). The figure highlights two subnetworks that we find particularly interesting. One is enriched in proteins that are connected to clathrin-coated vesicles (see below). The other is enriched in splicing factors (insets in Fig. 2 B) that interact with an IDR corresponding to amino acids 512–526 of Fused in sarcoma (FUS). These interactions are disrupted by the R521C mutant. FUS is an RNA-binding protein that is best known for its role in amyotrophic lateral sclerosis (ALS) ^30^. The R521C and other mutations in the C-terminal region of the protein are thought to be pathogenic because they disrupt a nuclear localisation signal ^31^. Our data suggests that impaired binding of splicing factors could be an additional/alternative explanation for the pathogenicity of this mutation. This observation is interesting because FUS has already been implicated in splicing ^32–34^. In fact, the C-terminal region of the protein was found to interact with SRSF10 even before pathogenic mutations in this region were identified ^35^.

## Recruitment of clathrins through gains of dileucine motifs

The finding we considered most interesting is that mutated IDRs from CACNA1H, GLUT1/SLC2A1 and ITPR1 led to specific interactions with clathrins (Fig. 3 A). The corresponding SILAC data revealed that in all three cases clathrin had a strong preference for the mutant form of the peptides over the wild-type (Fig. 3 B). Clathrins mediate endocytosis and intracellular trafficking of transmembrane proteins. They are recruited to the membrane by adaptor proteins that recognize specific cargo, form clathrin-coated pits which are then pinched off by dynamin ^36,37^. Our finding that clathrins are specifically recruited to mutated IDRs suggests that the mutations might affect protein trafficking. Intriguingly, the three mutations share several other features beyond an increased affinity for clathrin. First, all three mutations affect transmembrane proteins -- a calcium channel (CACNA1H) and a glucose transporter (GLUT1) residing in the plasma membrane and an inositol 1,4,5-trisphosphate receptor (ITPR1) in the ER (Fig. 3 C). Second, all three mutations affect disordered regions exposed to the cytosol, which makes them accessible to cytosolic adaptor proteins that mediate clathrin recruitment. Third, all three mutations involve the change of a proline to a leucine residue and thereby result in the appearance of a novel dileucine motif (“LL”) in the IDR (Fig. 3 D). Such motifs are known to recruit clathrin to the plasma membrane or intracellular locations ^38^. The classical dileucine motif for clathrin-dependent endocytosis is [D/E]XXXL[L/I] ^39^, but variations of this scheme are common ^38,40,41^.

**Fig. 3:**
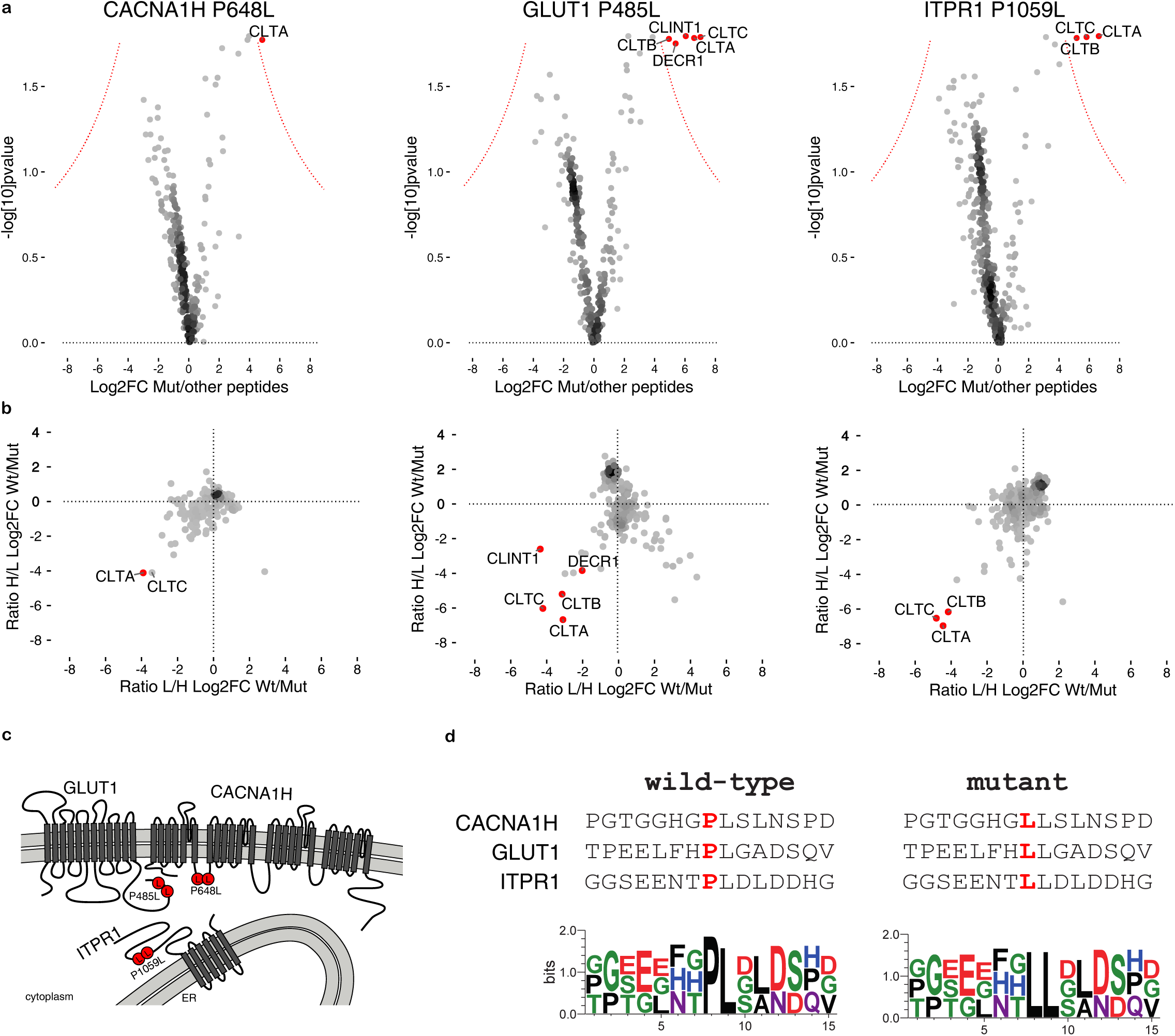
Recruitment of clathrins by recurrent gains of dileucine motifs. **A**, Volcano plots for pull-downs with mutated peptides derived from CACNA1H, GLUT1 and ITPR1. Specific binders (relative to all other pull-downs) are highlighted in red. All three peptides specifically interact with clathrins. **B**, Corresponding SILAC plots show that clathrins and related proteins preferentially bind to the mutant form of peptides (relative to the wild-type). **C**, Graphical representation of the mutation sites. All three mutations affect cytosolic regions of transmembrane proteins. CACNA1H and GLUT1 are located in the plasma membrane and ITPR1 in the ER. **D**, Aligning the three peptide sequences reveals a common gain of a dileucine motif.

## The P485L mutation causes mislocalization of the glucose transporter GLUT1

The data presented so far are derived from our artificial *in vitro* screen based on short peptides. We therefore selected the P485L mutation in GLUT1 for follow-up experiments. This mutation causes GLUT1 deficiency syndrome (G1DS), a disorder characterized by seizures and developmental delays ^42,43^. GLUT1 facilitates glucose transport into the brain across the blood-brain barrier. GLUT1 mutations in G1DS patients impair this glucose transport which gives rise to the disease phenotype.

To determine the impact of the P485L mutation on the subcellular localization, we generated stable inducible cell lines expressing epitope tagged full-length wild-type or mutant GLUT1. While the wild-type protein mainly localized to the plasma membrane, the P485L mutant showed a more vesicular pattern (Fig. 4 A). Next we tested whether the mutant form of GLUT 1 was taken up via endocytosis by adding fluorescently labeled transferrin to GLUT1 expressing cells. In contrast to wild-type GLUT1, the mutant extensively colocalized with endocytosed transferrin (Fig. 4 B). To systematically characterize the endocytic compartment in which GLUT1_P485L resides, we used BioID as a proximity labeling method ^44^. We performed this experiment in a comparative manner for both wild-type and mutant GLUT1 with SILAC-based quantification (see Methods). GLUT1_P485L showed increased colocalization with proteins involved in clathrin-mediated endocytosis, the trans-Golgi network (TGN), retrograde endosome-to-TGN transport and lysosomes (Fig. 4 C). Of note, AP2B1 also showed increased association with mutated GLUT1. This protein is part of the adaptor protein 2 (AP2) complex that recognizes dileucine motifs to mediate endocytosis ^38,40,41^. Together, these data indicate that the P485L mutation causes internalization of Glut1 via AP2-mediated clathrin-dependent endocytosis.

**Fig. 4:**
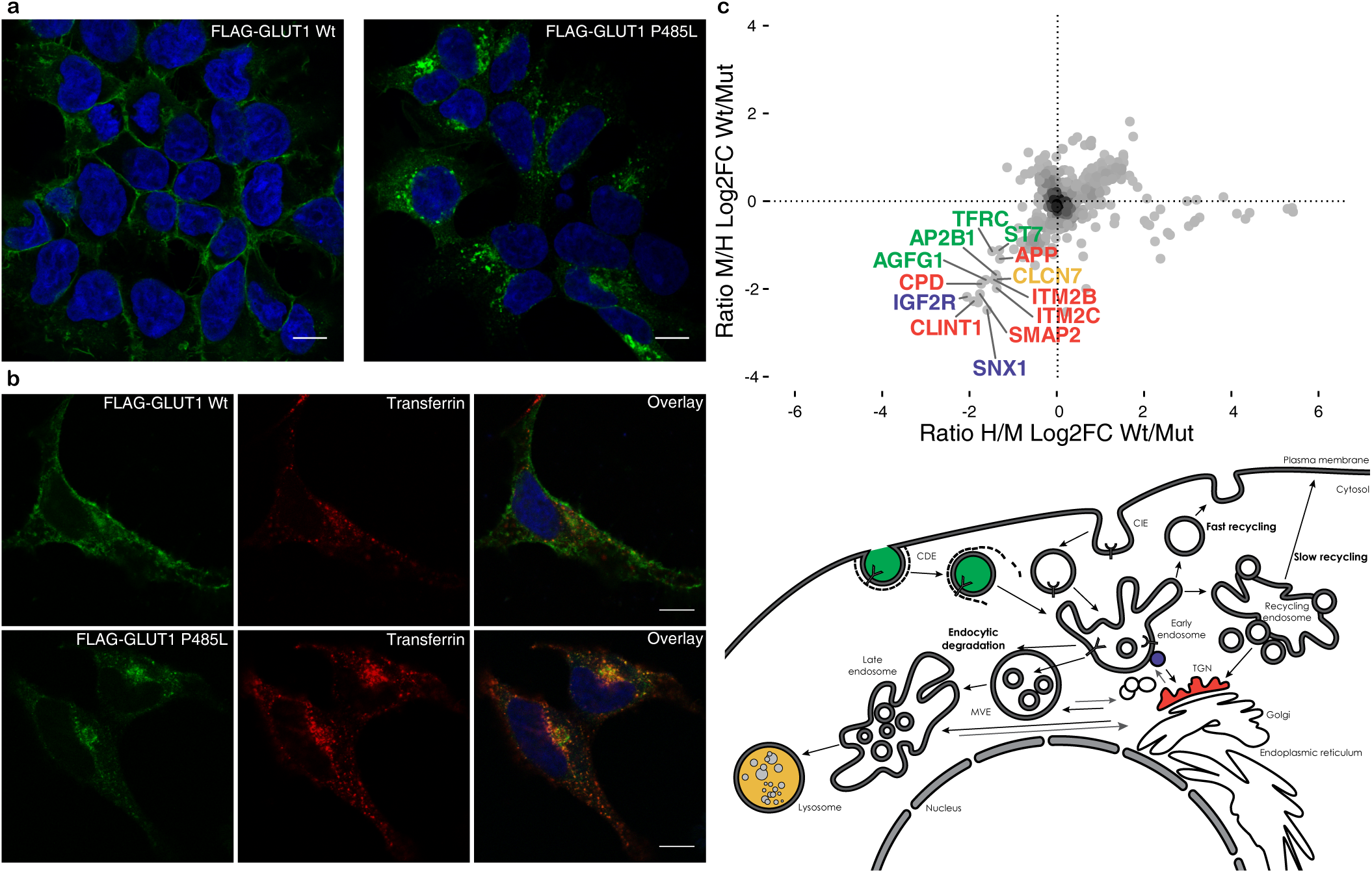
A mutation-induced dileucine motif gain causes mislocalization of the glucose transporter GLUT1. **A**, Confocal images of GLUT1 localization in Hek cells stably expressing FLAG-GLUT1 reveal that the wild-type is localized mainly at the cell membrane while the P485L mutant is mislocalized to cytoplasmic compartments. **B**, GLUT1 expressing HeLa cells are incubated with fluorescently labeled transferrin for 10 min before fixation. Mutant but not wild-type GLUT1 extensively co-localizes with endocytosed transferrin. **C**, Comparison of proteins co-localizing with wild-type and mutant GLUT1 by proximity labeling (BioID). The upper panel shows SILAC log2 fold changes from two replicate experiments with swapped isotope labels. Proteins are colored according to their typical subcellular compartment (lower panel). The P485L mutant shows increased colocalization with proteins involved in in clathrin-dependent endocytosis (CDE), the trans-Golgi network (TGN), retrograde endosome-to-TGN transport and lysosomes. Figure adapted from ^46^.

## Gains in dileucine motifs as a general disease mechanism

We next wondered if dileucine motif gains might be a more general mechanism by which diseases could arise. To test this, we conducted a search of missense mutations known to cause disease and occurring within disordered cytosolic regions of transmembrane proteins. We found four additional pathogenic dileucine motif gains (Fig. 5 A, Table S4). These mutations affect different proteins and cause a range of diseases. Since we focused our follow-up experiments on GLUT1, we cannot state with certainty if the other dileucine motif gains also cause protein mistrafficking. Alternatively, the mutations could cause disease by different mechanisms and might just create dileucine motifs as a by-product. If that was the case dileucine motifs should be homogeneously distributed between disease-causing mutations and non-pathogenic polymorphisms. Alternatively, if dileucine motif gains are responsible for pathogenesis, they would be predicted to occur more often in disease than in non-pathogenic variants. Moreover, pathogenic dileucine motif gains should be specific for cytosolic regions of transmembrane proteins since this is where they exert their function. To test these predictions, we compared the frequency of dileucine motif gains that have been found in disease-causing mutations to their appearance in non-pathogenic polymorphisms. A global survey of all disordered regions of the entire proteome revealed that gains in dileucine motif gains occurred at about the same rate in disease and non-pathogenic variants (OR = 0.81, p-value = 0.319, two-sided Fisher’s Exact Test). In the cytosolic tails of transmembrane proteins, however, we observed a 3.7-fold enrichment of dileucine motifs implicated in disease (OR = 3.7, p-value = 0.017, two-sided Fisher’s Exact Test, Fig. 5 B). Disordered extracellular regions of transmembrane proteins do not show this enrichment. We conclude that dileucine motif gains in disordered regions of the cytosolic segments of transmembrane proteins are significantly and specifically enriched in disease. To further assess the significance of this finding, we systematically searched within cytosolic regions of transmembrane proteins for all other annotated SLiMs contained in the ELM database ^39^. Intriguingly, of all 263 SLiMs tested, the dileucine motif (LIG_diLeu_1) was the only significantly enriched motif in disease (Fig. 5 C).

**Fig. 5:**
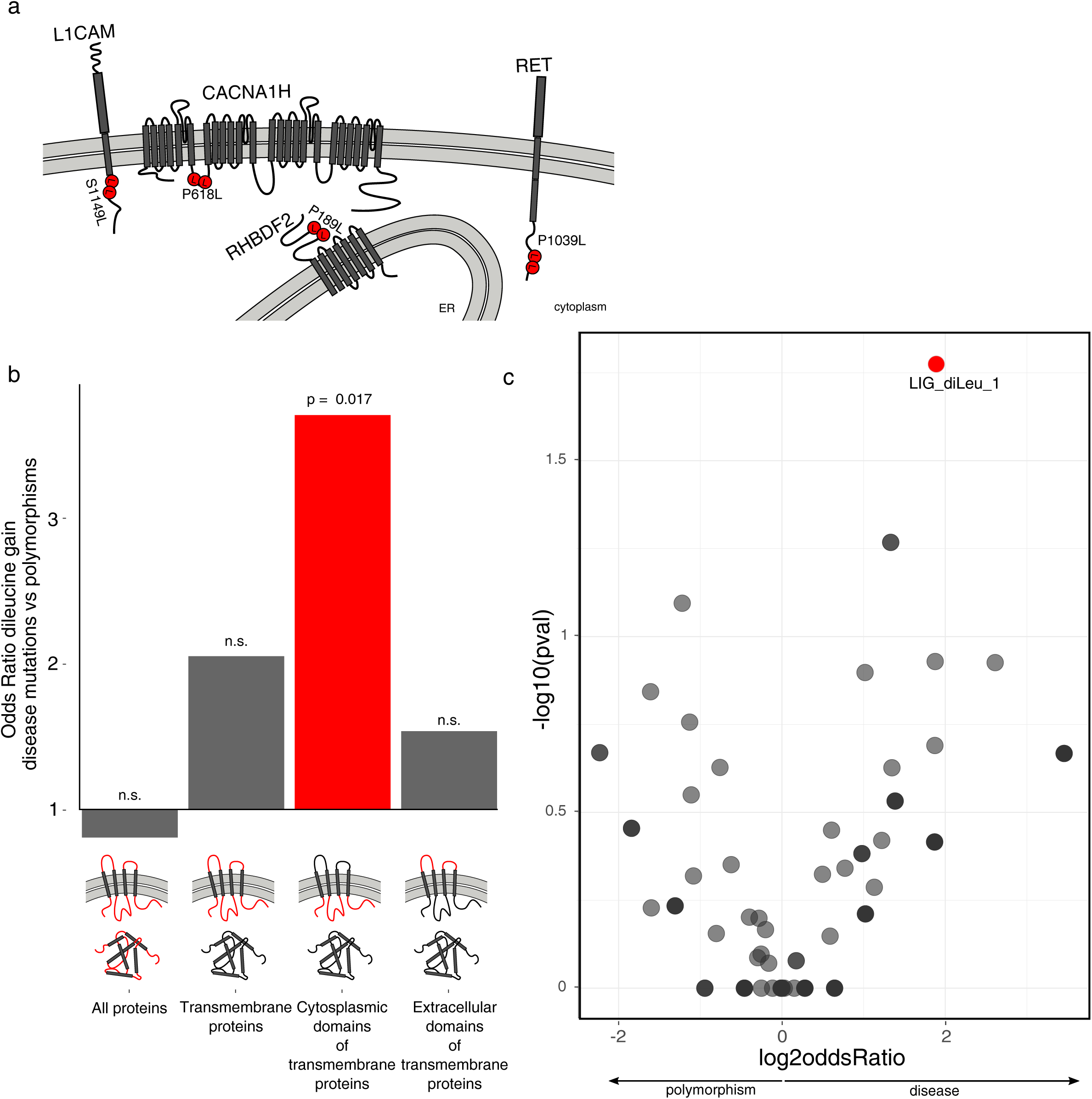
Mutation-induced gains of dileucine motifs are a significant cause of disease. **A**, A systematic bioinformatic search revealed four additional pathogenic mutations in cytosolic segments of transmembrane proteins that create dileucine motifs. **B**, Relative frequency of dileucine motif gains in disease mutations and polymorphisms in different disordered regions (IUPred Score >= 0.4) of the proteome. Dileucine motif gain is significantly enriched only in disordered regions of the cytoplasmic domains of transmembrane proteins (two-sided Fischer’s exact test). **C**, Comparison of gained motifs in disordered regions of cytoplasmic tails of transmembrane proteins reveals the dileucine motif to have the most significant specific enrichment when compared with polymorphisms.

## Discussion

Understanding the functional relevance of protein variants is a major challenge in the era of personal genomics -- especially for missense mutations in disordered regions. Our proteomic screen provides a first systematic experimental analysis of how mutations in disordered regions affect protein-protein interactions. Our results show that the method can (i) capture known interactions, (ii) detect how mutations in SLiMs affect binding of cognate domains and (iii) provide novel mechanistic insights into pathogenesis. The peptide-based method is especially useful for mutations in proteins that are otherwise difficult to study, such as large transmembrane proteins. Nevertheless, it is also important to consider the intrinsic limitations of the approach. Most importantly, *in vitro* pull-downs do not necessarily reflect physiological interactions *in vivo*. For example, artifactual binding can occur when combining peptides and proteins that never see each other in the cell ^45^. Moreover, taking IDRs out of the context of the full length protein and immobilizing them as short peptides can affect interactions. Finally, amino acids within IDRs often carry posttranslational modifications -- a possibility which we did not consider here. In the future, it will be interesting to include modified peptides, especially since mutations often affect modification sites ^19,20^.

Our screen revealed that a mutation in the cytosolic tail of the glucose transporter Glut1 generates a dileucine motif, which leads to clathrin-dependent endocytosis of the protein. Consistently, we found that mutant Glut1 mislocalizes from the plasma membrane to endocytic compartments. The finding that pathogenic mutations in cytosolic tails of other transmembrane proteins also create dileucine motifs is particularly intriguing. It is tempting to classify diseases caused by such motif gains as “dileucineopathies”. Whether the other dileucine motif gains cause protein mislocalization similar to Glut1 remains to be investigated. The observation that these mutations are significantly and specifically enriched in cytosolic tails suggests that at least some of them are functional. Why are dileucine motif gains a recurrent cause of disease? We think this is due to a combination of several factors: First, the dileucine motifs are not very complex and can thus arise by chance. Second, proline codons can mutate to leucine codons by changing a single base. Third, proline is overrepresented in IDRs, which are also the sites where the motif needs to be located in order to be functional.

Bioinformatic studies have established that pathogenic mutations in disordered regions often affect SLiMs ^19–21^. However, whether these predicted motif changes affect protein-protein interactions is not clear. Moreover, many motifs have not yet been defined and thus escape computational predictions ^14^. The biochemical approach presented here provides a useful complementary strategy to computational studies. Key advantages of our set-up are its scalability (by using synthetic peptides) and specificity (by employing two quantitative filters). While we focused on Mendelian neurological disorders here, the approach can also be applied to other types of variants such as somatic mutations in cancer and non-pathogenic polymorphisms.

## Acknowledgements

We would like to thank Michael Krauß, Volker Haucke (both FMP Berlin) and Philip Kim (University of Toronto) for helpful discussions and Russ Hodge (MDC Berlin) for valuable comments on the manuscript. We also thank Daniel Perez Hernandez (MDC Berlin) and Ulf Reimer (JPT) for practical suggestions in setting up the screen. Markus Landthaler (MDC Berlin) kindly provided stable cell lines and the Advanced Light Microscopy Technology Platform (MDC Berlin) helped with microscopy.

## Author contributions

K.M. and M.K. established the screen and selected the candidates. K.M. performed most of the wet-lab experiments with contribution from M.K. and J.C.. K.M. processed and analyzed the mass spectrometric data. B.U. carried out most of the remaining bioinformatic analyses (mainly motif based analyses) supervised by A.A.. M.S. conceived and supervised the work and wrote the manuscript with input from all authors.

## Materials and Methods

### Peptide-protein interaction screen

#### Candidate selection

Disease mutations in humans were taken from UniProt annotations ^1^ of Online Mendelian Inheritance in Man, OMIM^®^. McKusick-Nathans Institute of Genetic Medicine, Johns Hopkins University (Baltimore, MD), https://omim.org/. This dataset consists of experimentally validated missense mutations that contribute to inherited diseases. Inherited disease mutations were downloaded from UniProt (http://www.uniprot.org/docs/humsavar.txt, Release: 2015_07 of 24-Jun-2015, ^2^). Only mutations that were associated to ‘Disease’ were kept. ‘Unclassified’ mutations or ‘Polymorphisms’ were excluded. The 26,649 disease mutations were further filtered by applying a disorder cut-off. Disorder tendencies of 15 amino acids (AAs) long peptides, with the AA mutated in disease if possible located at position eight, were predicted using lUPred^3^ using the ‘SHORT’ profile considering sequential neighborhood of 25 residues. IUPred disorder scores above 0.5 denote regions of the proteins that have 95% likelihood to be disordered. For filtering, the mean disorder score for all 15AA as well as the mutation position were required to be >0.5. This resulted in 1,878 disease mutations in disordered regions. Next we assigned disease classes to 3,119 different diseases included in the humsavar database by combining a manual approach with automatic annotation with the Human Phenotype Ontology database, HPO ^4^. We selected 305 mutations causing neurological diseases. After manual inspection, we remained with 128 mutations causing 124 distinct neurological diseases that were used for the peptide-protein interaction screen.

#### Experimental setup

Peptides of 15 AAs, in total 128 wild-type peptide and 128 related peptides containing the disease causing mutation (256 peptides) plus one control peptide pair were synthesized in situ on cellulose membrane using PepTrack™ techniques (JPT Peptide Technologies, Berlin, Germany). Two of those peptide filters were moistened in cell lysis buffer [50 mM HEPES pH 7.6 at 4°C, 150mM NaCl, 1 mM EGTA, 1mM MgCl2, 10% Glycerol, 0.5% Nonidet P-40, 0.05% SDS and 0.25% Sodiumdeoxycholate, supplemented with protease inhibitor (Roche) and benzonase (Merck)]. In order to reduce unspecific binding, the membrane was incubated with 1 mg/ml yeast t-RNA (Invitrogen) for 10 min and then washed twice with cell lysis buffer. The entire peptide libraries were incubated with 15 ml of light or heavy SILAC labeled cell lysate (5 mg/ml) from SH-SY5Y cells for 2h. Membranes were washed three times and air dried.

#### Cell culture

SH-SY5Y, Hek293 and HeLa cells were cultured under standard cell culture conditions. In brief, cells were cultured in DMEM (life technologies) complemented with 10% fetal calf serum (Pan-Biotech).

Cells used for SILAC based experiments were cultured in SILAC DMEM (life technologies) complemented with glutamine (Glutamax, life technologies), Pyruvate (life technologies), non-essential amino acids (life technologies) and 10% dialyzed fetal calf serum (Pan-Biotech). The SILAC DMEM was supplemented with standard L-arginine (Arg0, Sigma-Aldrich) and L-lysine (Lys0, Sigma-Aldrich)(light”) as in ^5^. Alternatively, Arg6 and Lys4 (“medium-heavy”) or Arg10 and Lys8 (“heavy”) were added in place of their light counterparts. Cells were cultured at 37°C and 5% CO_2_.

#### Sample preparation for mass spectrometric analysis

Single spots were punched out from cellulose membrane with a 2mm diameter ear punch (Carl Roth) and SILAC pairs were placed together in a 96-well plate (Thermo Scientific) prepared with 30 μl of denaturation buffer [6M urea (Sigma-Aldrich), 2M thiourea (Sigma-Aldrich), 10mM HEPES, pH 8]. Samples were reduced by incubating with 10μl of 3.3 mM DTT (Sigma-Aldrich) for 30 min at RT, followed by an alkylation step using 10μl of 18.3 mM iodoacetamide (IAA) (Sigma-Aldrich) for 60 min at RT. The samples were first digested using 1μg endopeptidase LysC (Wako, Osaka, Japan) for 4 hours. The samples were diluted by adding 100 μl of 50mM ammonium bicarbonate (pH = 8.5), and finally digested with 1μg trypsin (Promega) for 16h. The digestion was stopped by acidifying each sample to pH < 2.5 by adding 10% trifluoroacetic acid solution. The peptide extracts were purified and stored on stage tips according to ^6^.

#### LC-MS/MS analysis

Peptides were eluted using Buffer B (80% Acetonitrile and 0.1% formic acid) and organic solvent was evaporated using a speedvac (Eppendorf). Samples were diluted in Buffer A (5% acetonitrile and 0.1% formic acid). Peptides were separated on a reversed-phase column with 45 min gradient with a 250 nl/min flow rate of increasing Buffer B concentration on a High Performance Liquid Chromatography (HPLC) system (ThermoScientific). Peptides were ionized using an electrospray ionization (ESI) source (ThermoScientific) and analyzed on a Q-exactive plus Orbitrap instrument (ThermoScientific). Dynamic exclusion for selected precursor ions was 30 s. The mass spectrometer was run in data dependent mode selecting the top 10 most intense ions in the MS full scans, selecting ions from 300 to 1700 m/z (Orbitrap resolution: 70,000; target value: 1,000,000 ions; maximum injection time of 120 ms). The resulting MS/MS spectra from the Orbitrap had a resolution of 17,500 after a maximum ion collection time of 60 ms with a target of reaching 100,000 ions.

#### Data analysis

The resulting raw files were analyzed using MaxQuant software version 1.5.2.8 ^7^. Default settings were kept except that ‘match between runs’ and ‘re-quantify’ was turned on. Lys8 and Arg10 were set as labels and oxidation of methionines and n-terminal acetylation were defined as variable modifications. Carbamidomethyl of cysteines was set as fixed modification. The in silico digests of the human Uniprot database (2015–12), a fasta file containing all peptides used for pull-down and a database containing common contaminants were done with Trypsin/P. The false discovery rate was set to 1% at both the peptide and protein level and was assessed by in parallel searching a database containing the reversed sequences from the Uniprot database. Following statistics and figures were done using R (R version 3.2.1, RStudio Version 1.0.143).

The resulting text files were filtered to exclude reverse database hits, potential contaminants, and proteins only identified by site. Missing LFQ-intensity values were imputed with random noise simulating the detection limit of the mass spectrometer ^8^. To this end, imputed values were taken from a log normal distribution with 0.25× the standard deviation of the measured, logarithmized values, down-shifted by 1.8 standard deviations. In this way, a distribution of quantitative values for each protein across samples is obtained. For determination of specific interactions, two replicated pull-downs for the same peptide were tested against all other pull-downs, excluding the corresponding variant peptide, by the nonparametric Mann–Whitney U test. Resulting p-values and fold-changes (log2 space) have been plotted as volcano plots to determine cut-offs. For cut-offs, an approach was used that employs a graphical formula to combination a fold-change and p-value cut-off ^8^: 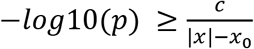with x: enrichment factor of a protein, p: p-value of the Mann–Whitney U test calculated from replicates, x_0_: fixed minimum enrichment, c: curvature parameter. The curvature parameter c determines the maximum acceptable p-value for a given enrichment x.

The parameters c and x_0_ can be optimized based on prior knowledge of known true and false positives ^8^. Here, cut-offs were chosen according to known interaction partners of the SOS1 control peptide ^8,9^. This resulted in x_0_=0, c=8.

This cut-off was applied to all other pull-downs to separate specific binders from background. SILAC ratios were normalized by subtracting the median SILAC ratio of every experiment from all SILAC ratios in that experiment. To define interaction partners that bind differentially to wild-type and mutant peptide, a SILAC cut-off has been defined. For wild-type specific interaction partners, the mean SILAC ratio of the two replicates needed to be >1 and none of the two ratios <0 (mutant specific mean SILAC ratio < -1 and none of the two ratios >0). Resulting figures were modified in Inkscape (0.91).

### Follow-up on GLUT1

#### Generation of vectors

We purchased SLC2A1 (GLUT1) from Harvard Plasmid repository (HsCD00378964). A stop codon has been added to the gene with the following primers Fw:TCCCAAGTGTAATTGCCAACTTTCTTGTACAAAGTTG, Rev:ATCAGCCCCCAGGGGATG. P485L Mutation has been introduced by changing c.1454 C>T ^10^ with Q5^®^ Site-Directed Mutagenesis Kit (NEB) Fw:CTGTTCCATCtCCTGGGGGCT, Rev:CTCCTCGGGTGTCTTGTCAC. SLC2A1 has been further cloned into two destination vectors adding either an N-terminal FLAG-HA Tag or a BirA-FLAG Tag with Gateway cloning strategy (Thermo Fisher Scientific).

#### BioID

Light, medium-heavy and heavy labelled Hek293 cells have either been mock-transfected (light cells served as a control for background binding) or transiently transfected with BirA-FLAG-GLUT1 wt or mutant (medium-heavy and heavy labeled cells). SILAC labeling allowed for quantitative comparison of proteins that have been proximity labelled by the transiently expressed constructs (Forward experiment: Light - Control, Medium-heavy - wt, Heavy - mut; Label swap experiment: Light - Control, Medium-heavy - mut, Heavy - wt). All three cell lines have been incubated for 24h in cell culture medium containing biotin. BioID experiment has been performed essentially as in ^11^, with minor adaptations.

Mass spec setup and analysis was done similarly as to samples from peptide pull-downs, but digested peptides were separated on a 2,000 mm monolithic column with a 100-μm inner diameter filled with C18 material that was kindly provided by Yasushi Ishihama (Kyoto University) using a 4 hr linear gradient with a 300 nl/min flow rate of increasing Buffer B concentration on a High Performance Liquid Chromatography (HPLC) system (ThermoScientific).

#### GLUT1 localization

Hek293 cells stably expressing either FLAG-tagged GLUT1 wt or P485L mutant exhibiting tetracycline-inducible expression were generated using the Flp-In system developed by Life Technologies according to the manufacturer’s protocol. After induction for 24h in doxycyclin (0.1 μg/ml) containing media, cells have been stained against FLAG (F1804, SIGMA). Nucleus has been stained with DAPI (Sigma).

#### Transferrin uptake

HeLa cells seeded on coverslips coated with poly-l-lysine (Sigma) have been transiently transfected with FLAG-GLUT1-wt or -P485L construct. After 24h they were serum-starved for 1 h and used for Transferrin (Tf) uptake. For Tf uptake, cells were treated with 20 μg ml^−1^ Tf-Alexa568 (life technologies) for 10 min at 37 °C.

#### Fluorescence Microscopy

Cells were cultured on poly-l-lysine (Sigma) coated coverslips and fixed with 4% PFA. Standard procedures were used for immunostaining. Images were acquired by confocal microscopy (Transferrin uptake: Zeiss Observer.Z1; GLUT1 localization: Leica, DMI6000). Images were further processed with Fiji ^12^.

### Analysis of human missense variants and short linear motifs (SLiMs)

#### SLiM regular expression patterns

262 annotated SLiM class definitions (regular expression patterns) were downloaded from the Eukaryotic Linear Motif (ELM) database ^13^. In order to analyse dileucine motifs, an additional motif ‘.LL.’ was added to this compilation and named ‘LIG_diLeu_1’ in order to conserve the naming convention followed by the ELM database.

#### Pathogenic and non-pathogenic missense variants

For the analysis of the missense variants that lead to *de novo* SLiM instances in protein sequences **Uniprot Humsavar dataset** (version 12-Apr-2017) ^2^ was downloaded and filtered for missense variants. Variants that are classified as ‘Disease’ or ‘Polymorphism’ in this dataset were selected.

#### Protein domains

PFAM domain annotations of proteins were downloaded from the PFAM database (ftp://ftp.ebi.ac.uk/pub/databases/Pfam//releases/Pfam30.0/proteomes/9606.tsv.gz) ^14^.

#### SLiM - PFAM associations

PFAM domains and SLiM classes that are known to interact were downloaded from the ELM database (http://elm.eu.org/interactiondomains).

#### Analysis of gain of SLiMs via missense variants in disordered regions

For each reviewed human protein from Uniprot (20191 proteins), the disorder scores of each residue were calculated using IUPred (using the ‘short’ setting). Using a IUPred disorder score cut-off of 0.4, the missense variants in disordered regions were selected. The missense variants that overlap PFAM domains were further filtered out based on the PFAM domain annotations found in the protein feature files downloaded from Uniprot in GFF format (e.g. the link to the GFF file for GLUT1 is http://www.uniprot.org/uniprot/P11166.gff). These protein feature files were also used to detect the transmembrane proteins and their cytoplasmic/extracellular regions. The missense variants in disordered regions and not overlapping any PFAM domains were further classified as variants from 1) the whole proteome, 2) the transmembrane proteins (only those that have annotation of at least one cytoplasmic domain or an extracellular domain, in total 3836 proteins), 3) the cytoplasmic domains of transmembrane proteins, and 4) extracellular domains of transmembrane proteins. For each of these classes, the number of disease-causing variants and the number of polymorphisms that lead to a gain of SLiMs was counted and a two-sided Fisher’s Exact Test was applied to see if there is a statistically significant difference for the likelihood of a given class of SLiMs to be gained via disease-causing variant compared to that of polymorphisms.

#### Peptide-Protein Interaction Network Analysis

180 peptide-protein interactions that passed the strict LFQ filter and showed significant differential SILAC ratios between wild-type and mutant forms of the peptides were used to compose a peptide-protein interaction network. The network was visualized using Cytoscape 3.5.1 ^15^. Sub-graphs of the significant interactions were generated using R package igraph (version 1.0.1) ^16^ (using *fastgreedy.community* function) and visualized using the R packages ggnetwork ^17^ and ggplot2 ^18^. Enriched GO terms for each subgraph were calculated using the topGO R package ^19^.

## Supplementary figure legends

**Fig. S1:**
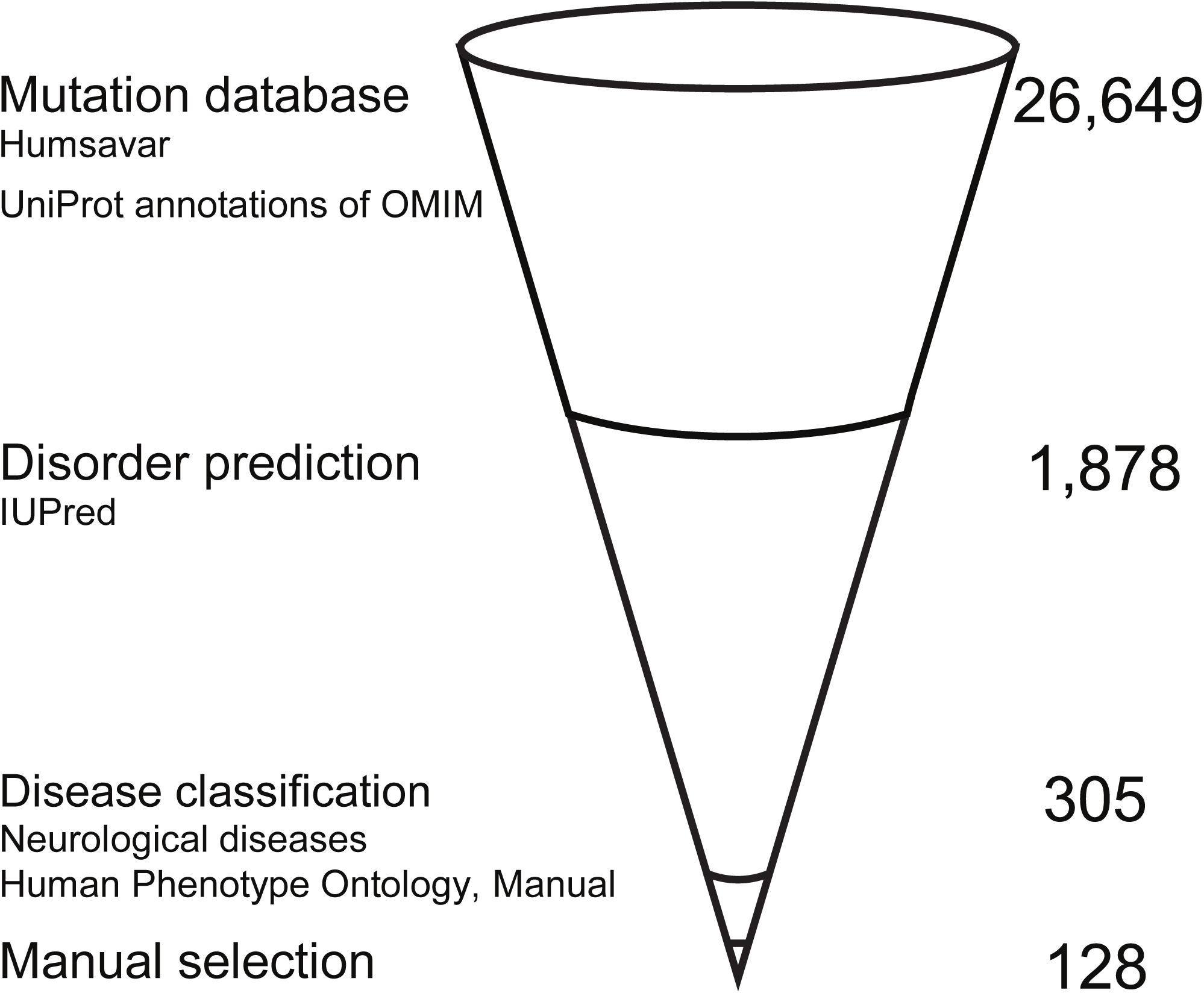
Candidate selection for peptide-protein interaction screen. Candidates were selected from missense disease mutations in the Humsavar database (Uniprot) by selecting mutations in disordered regions that cause neurological diseases. Only mutations in disordered regions of a protein were kept. Mutations were further filtered for those that cause neurological diseases and the final candidate set was manually selected.

**Fig. S2:**
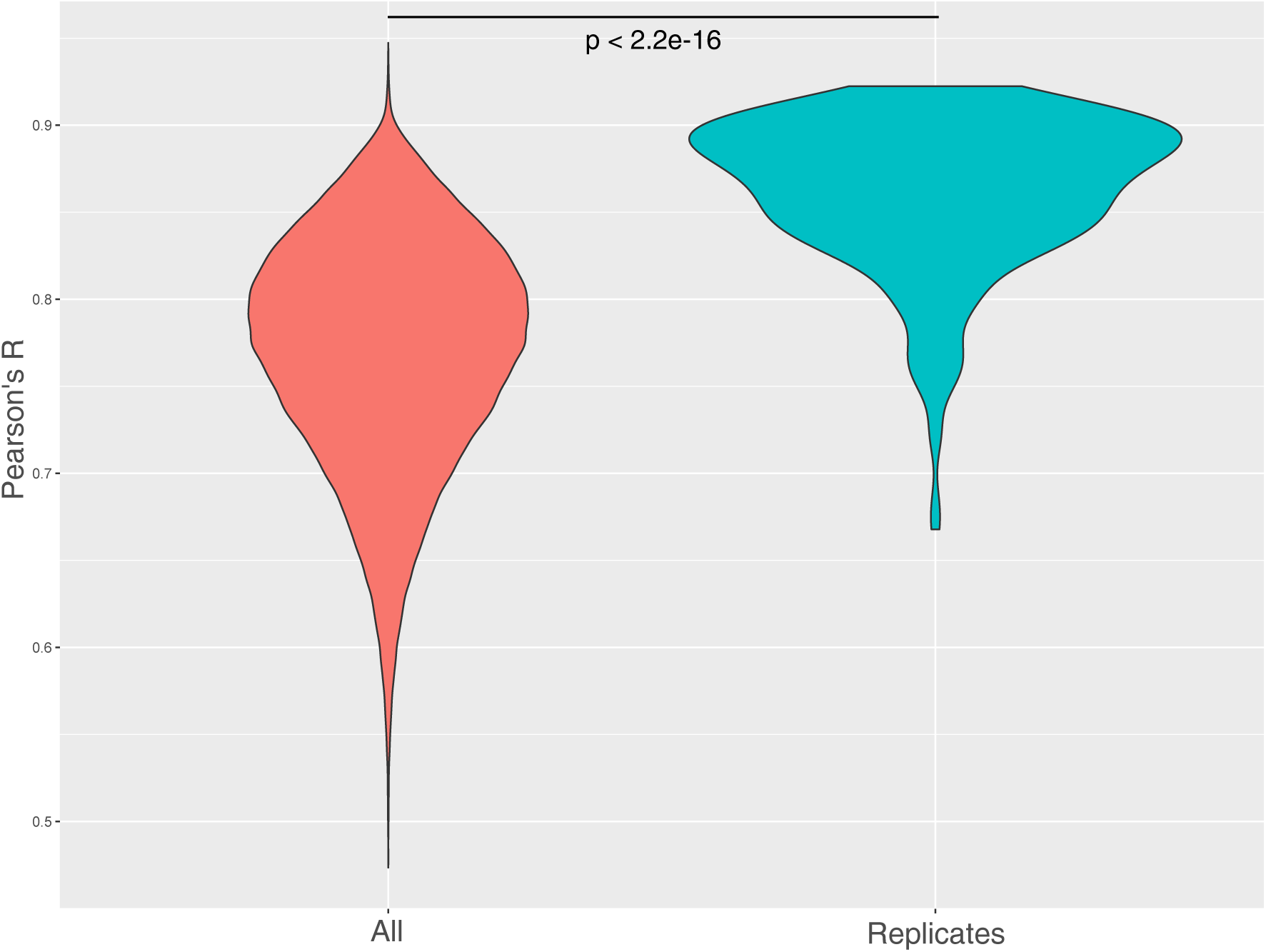
Reproducibility of technical replicates. Pearson’s R shows significantly higher correlation of technical replicates than correlations between all pull-downs.

**Fig. S3:**
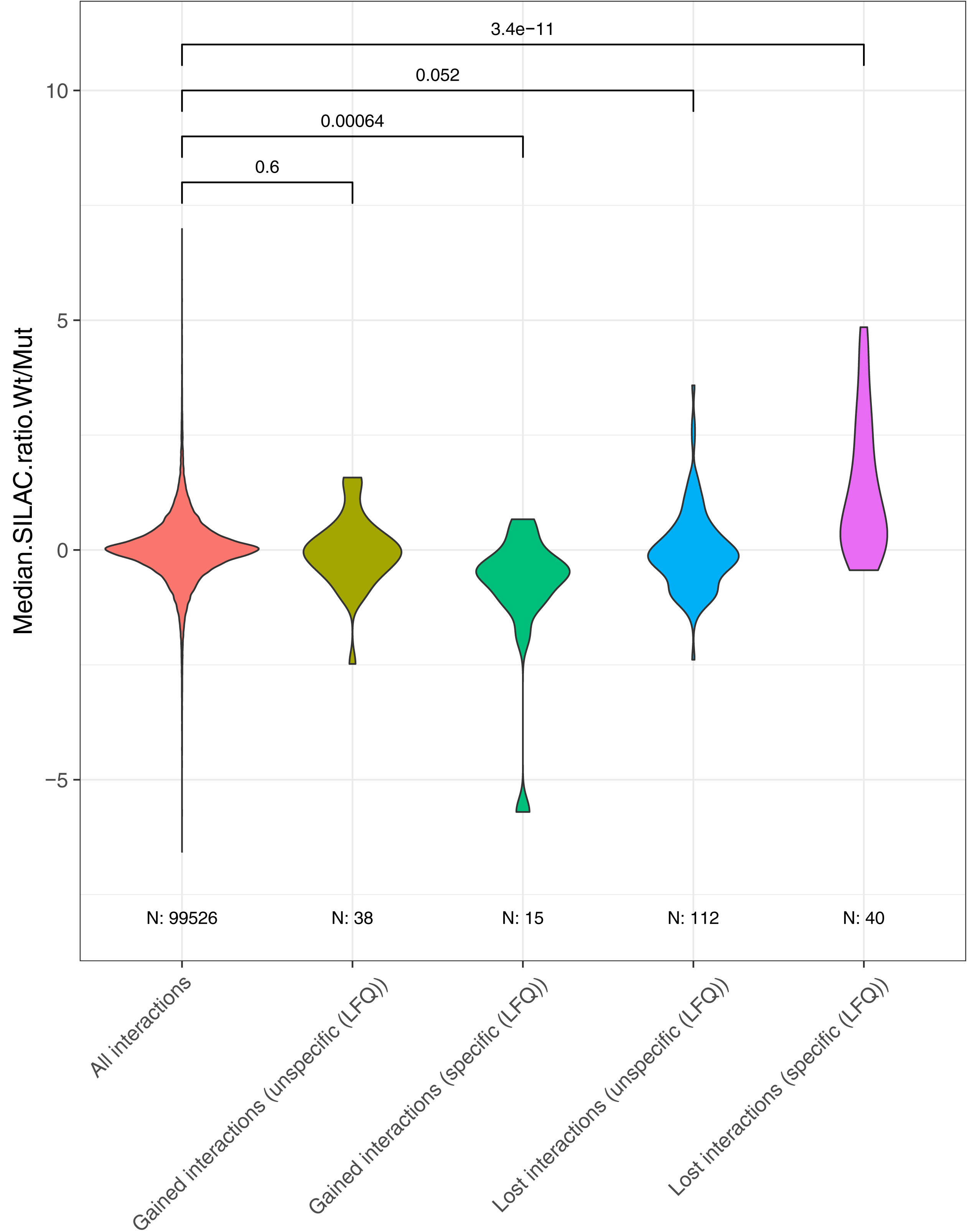
SILAC ratio distributions of detected interactions that can be explained by presence of SLiMs in the peptides and PFAM domains in the interaction partners. Peptide-protein interactions detected in the screen were classified as ‘gained’ or ‘lost’ according to the following criteria: An interaction is classified as ‘gained’ if the mutant peptide sequence matches a SLiM pattern that doesn’t match the wild-type peptide sequence and the mutant peptide has an interaction partner that contains a compatible PFAM domain to bind that SLiM instance. On the other hand, an interaction is classified as ‘lost’ if the wild-type peptide sequence matches a SLiM pattern that is not matched in the mutant peptide sequence and the wild-type peptide has an interaction partner that contains a compatible PFAM domain to bind that SLiM instance. Gained and lost interactions are further sub-classified as ‘LFQ positive’ and ‘LFQ negative’ depending on whether the peptide-protein interaction has an LFQ value that passes the loose LFQ cut-off. The median SILAC ratio distributions (wild-type versus mutant) of each of these four categories of interactions (‘Gained interactions - LFQ negative’, ‘Gained interactions - LFQ positive’, ‘Lost interactions - LFQ negative’, and ‘Lost interactions - LFQ positive’) are compared with the median SILAC ratio distributions of all detected interactions from the array using a Wilcoxon-Mann-Whitney test. Compared to the background distribution of median SILAC ratios (in red), the gained interactions that pass the LFQ filter (in green) show a significant negative skew while the lost interactions that pass the LFQ filter (in purple) show a significant positive skew.

**Fig. S4:**
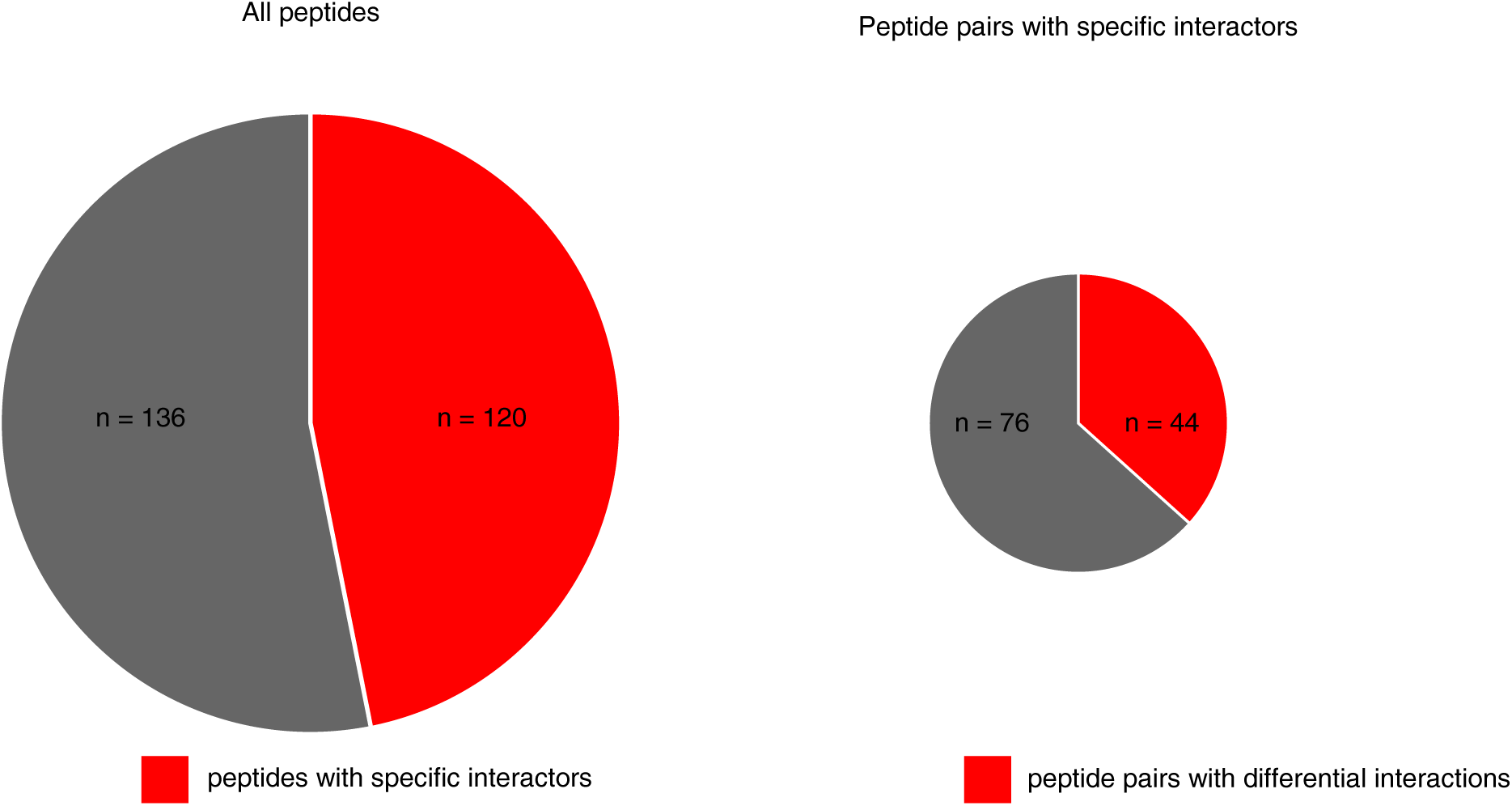
Impact of specificity cut-off (LFQ) and differential cut-off (SILAC) on peptide candidates. After applying the specificity cut-off (derived from control peptide, see Methods) on all interactions, only about half of the 2×128 peptides showed at least one specific binder according to the LFQ-filter (left pie chart, red). More than one third of all peptide pairs with specific interactors for wild-type and/or mutant show differential interactions after applying the SILAC cut-off (see Methods) (right pie chart, red).

**Fig. S5:**
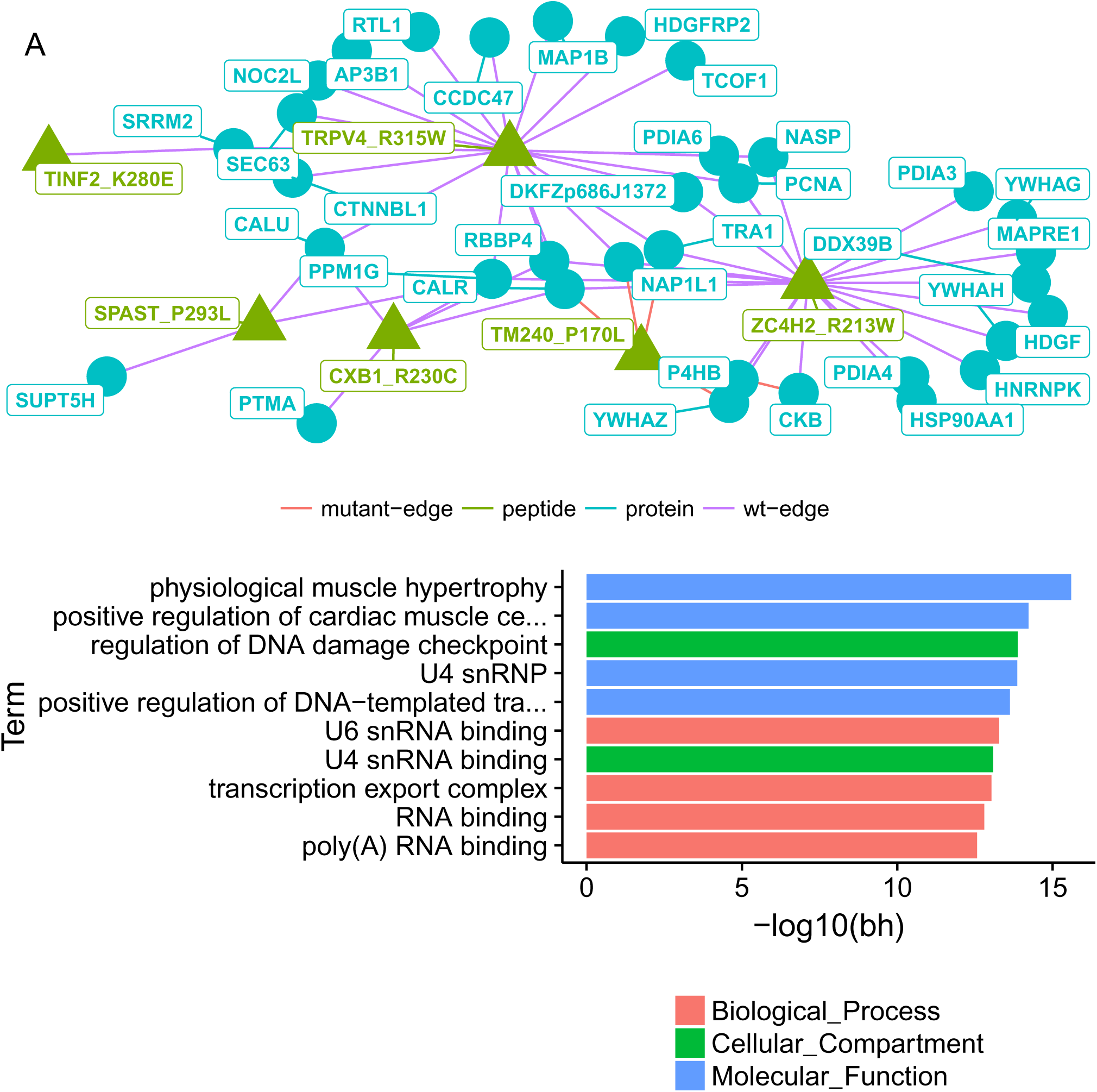
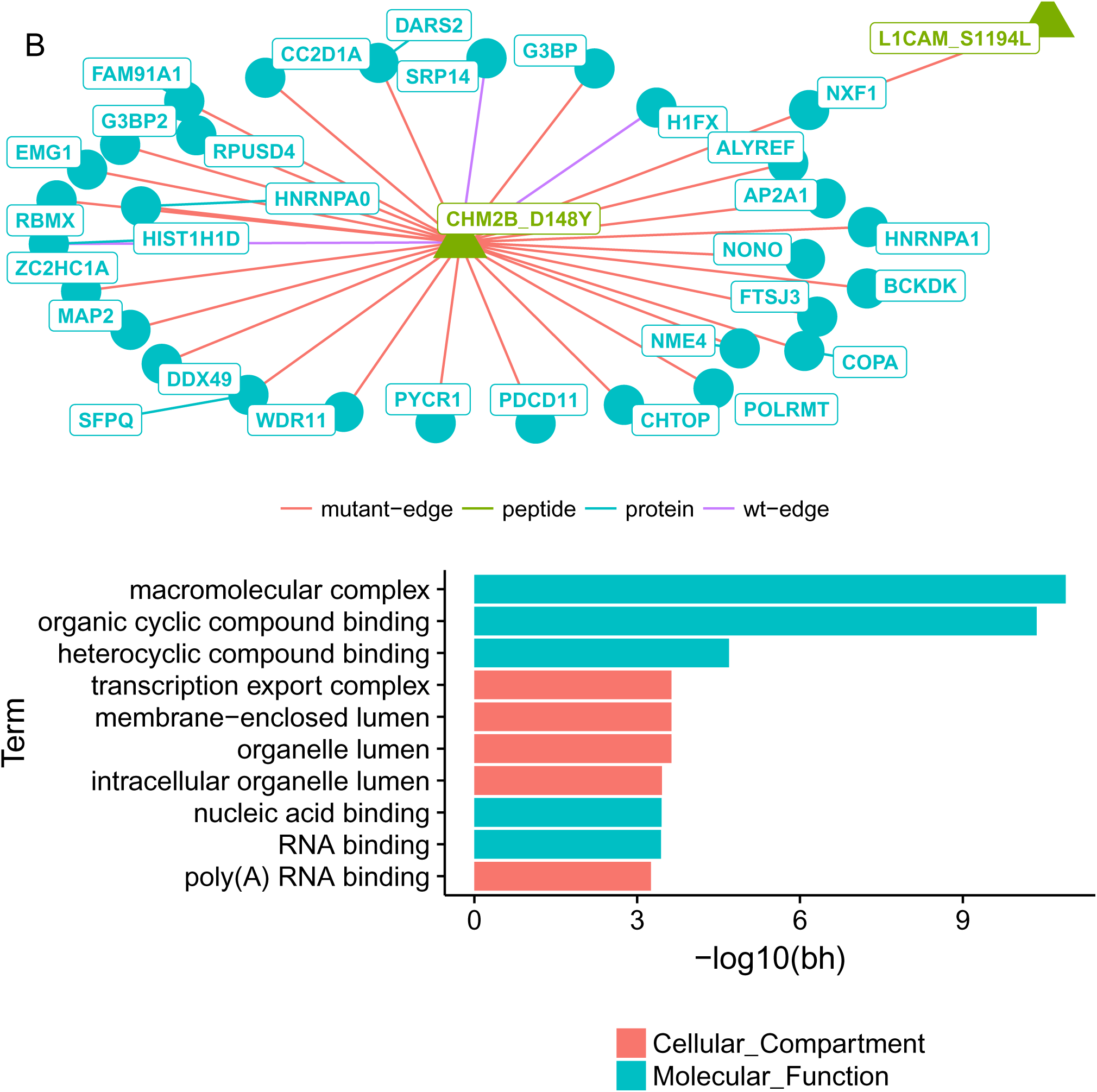
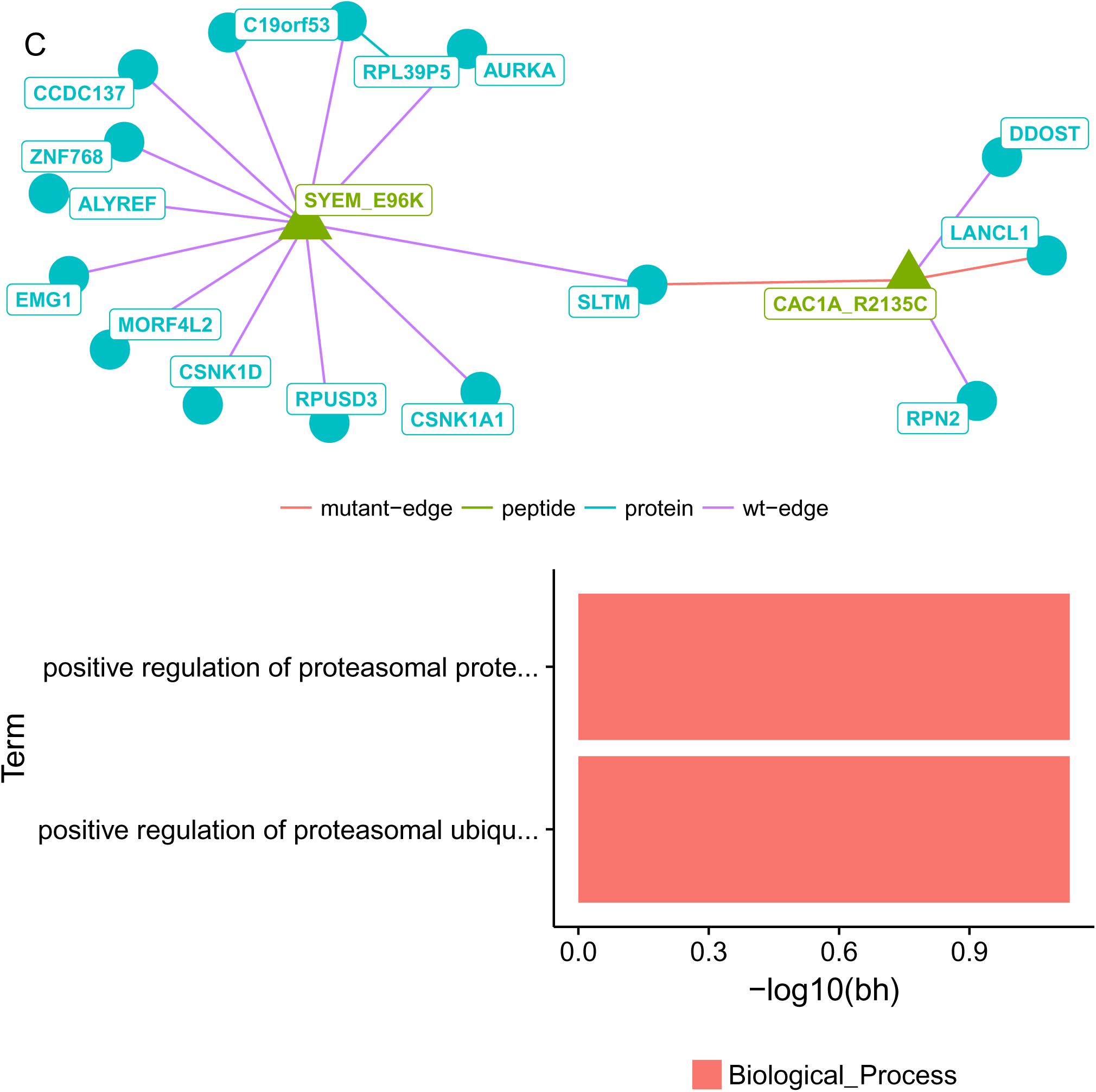
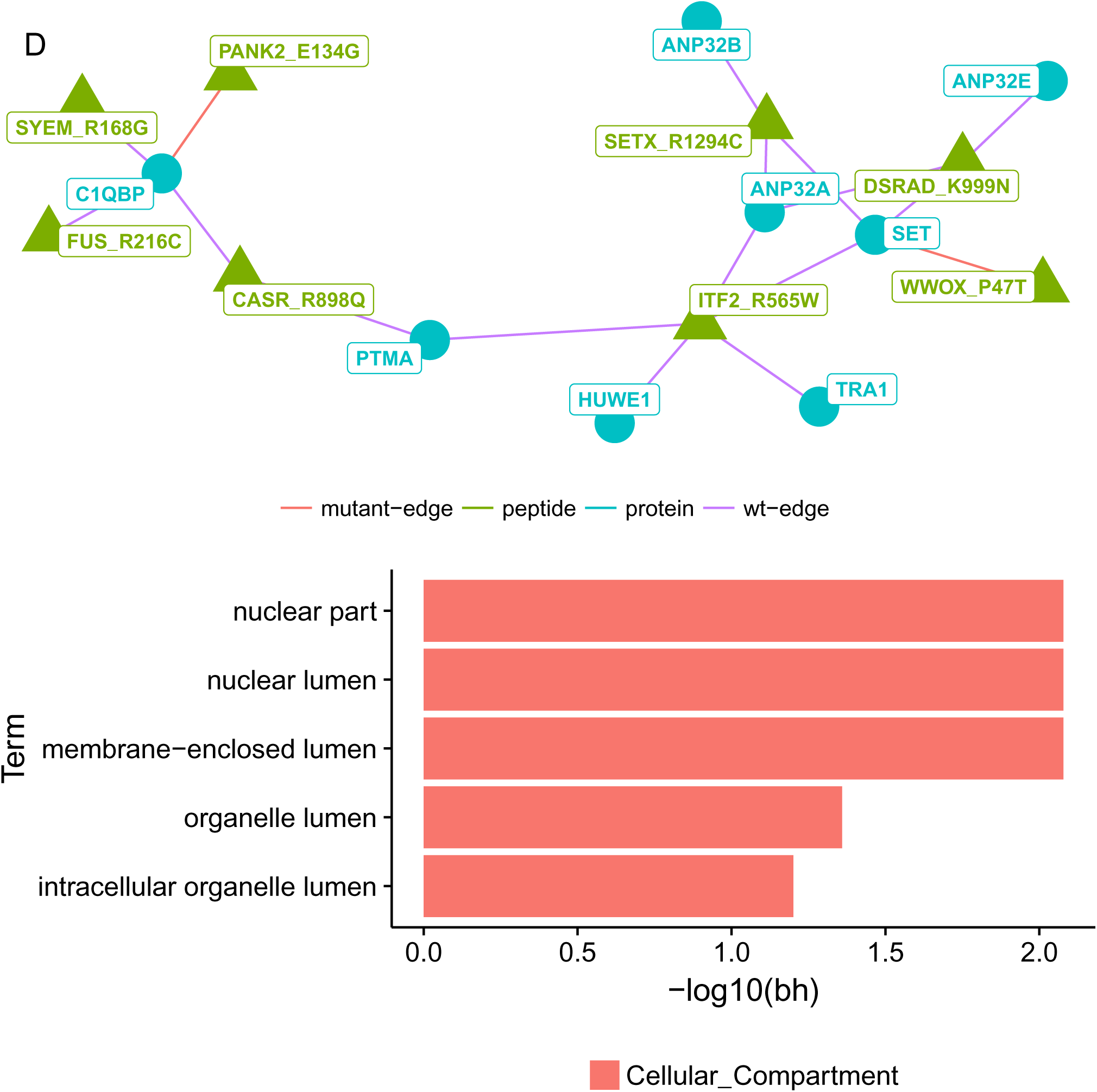
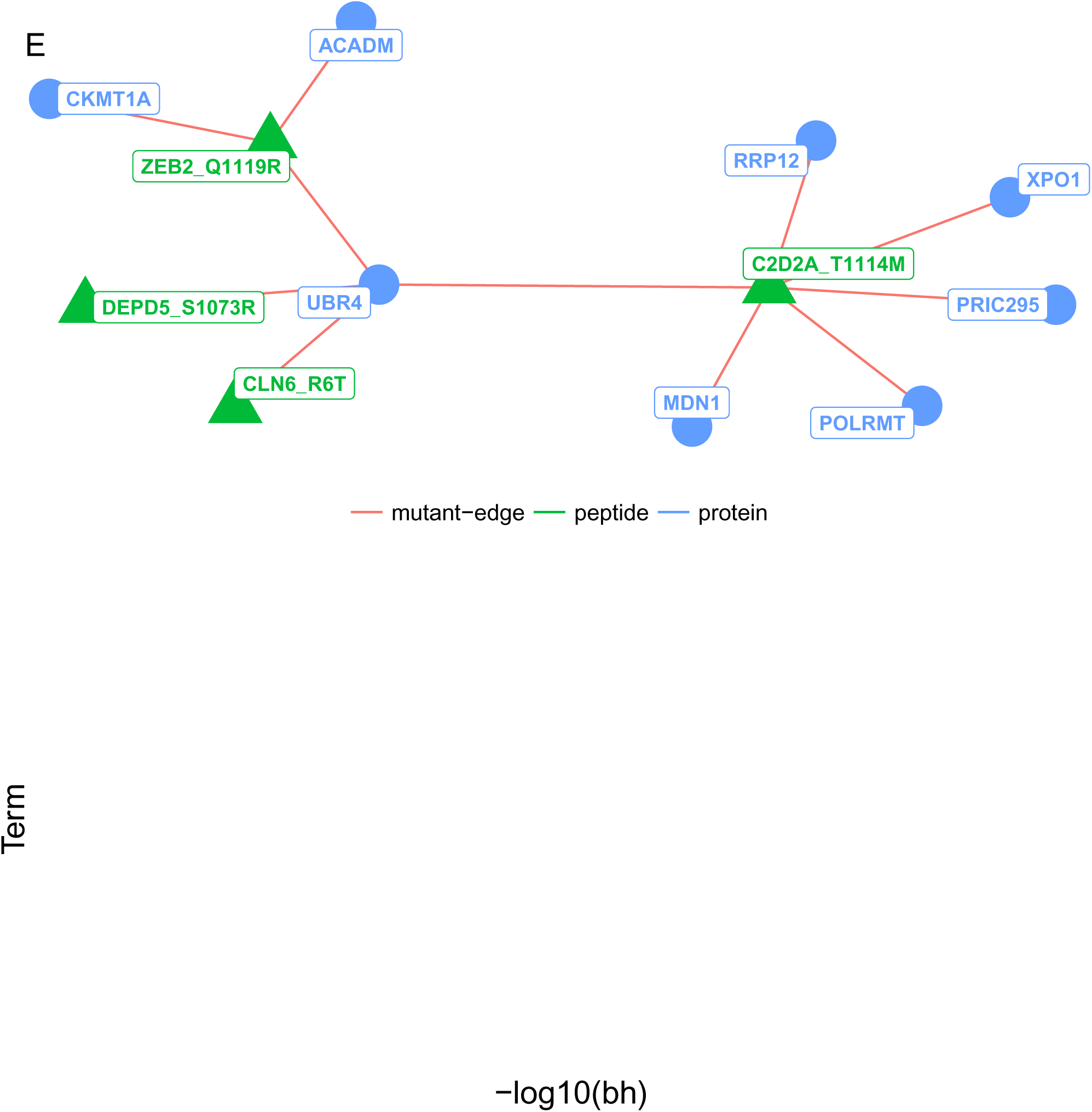
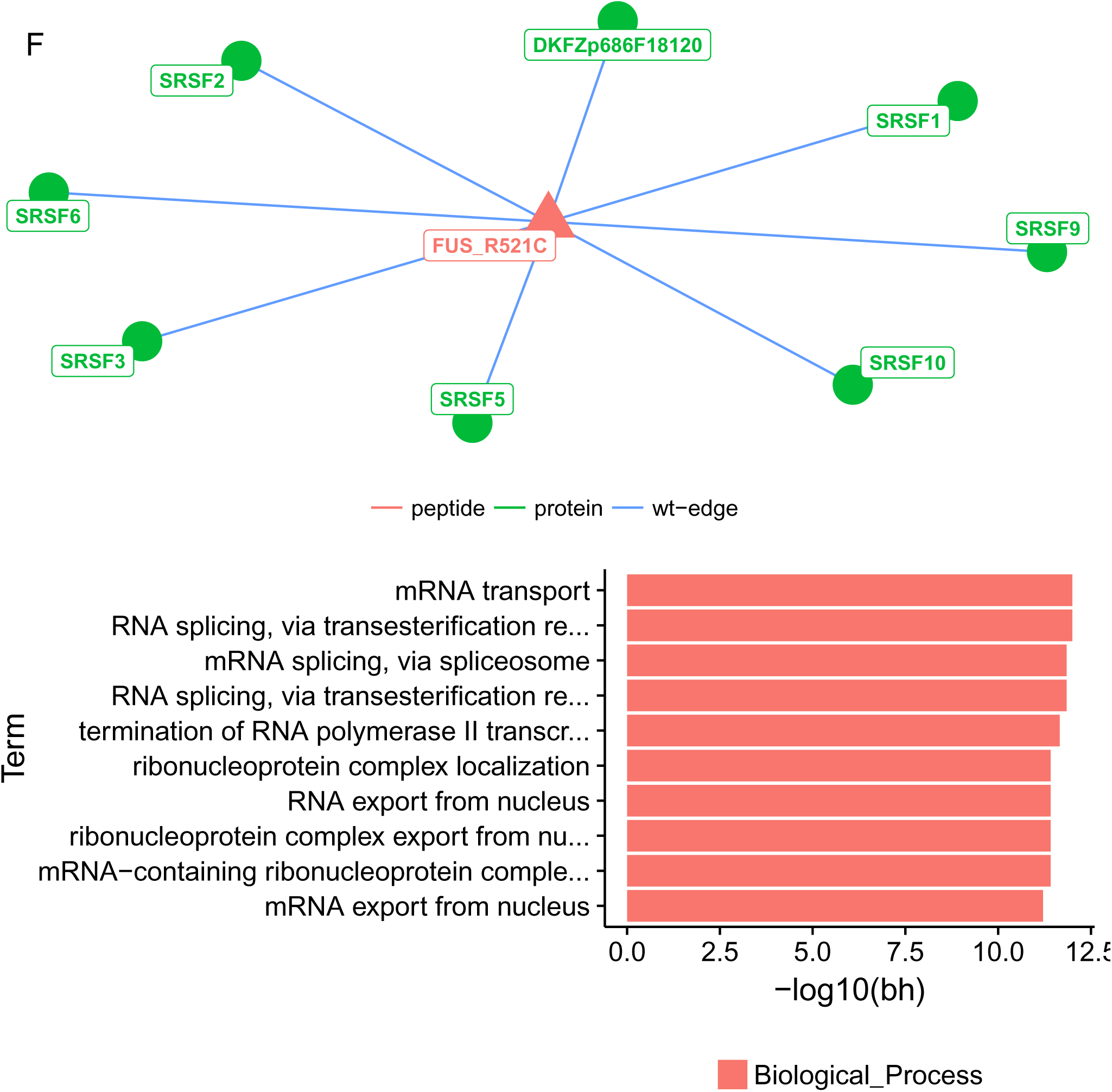
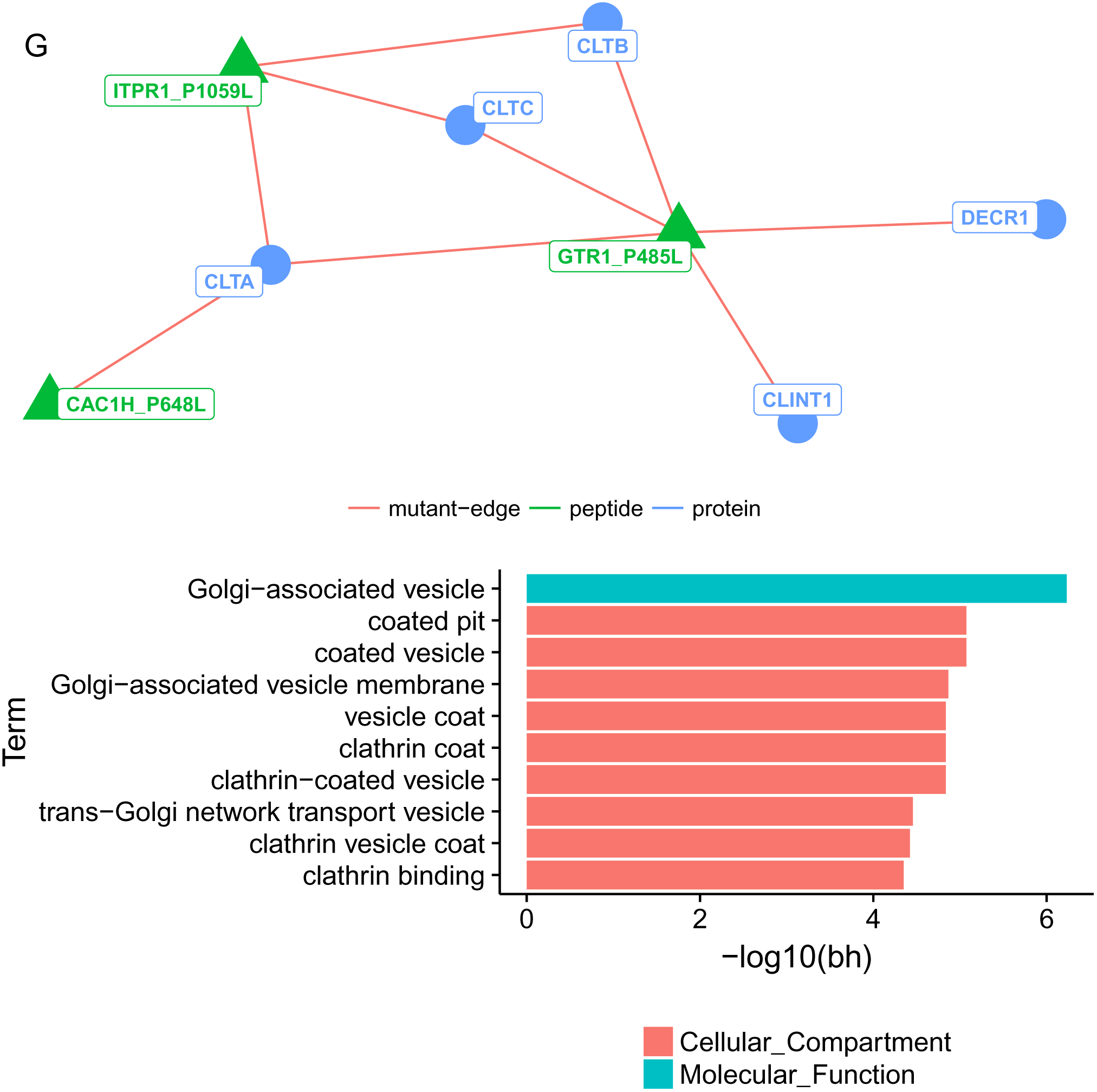
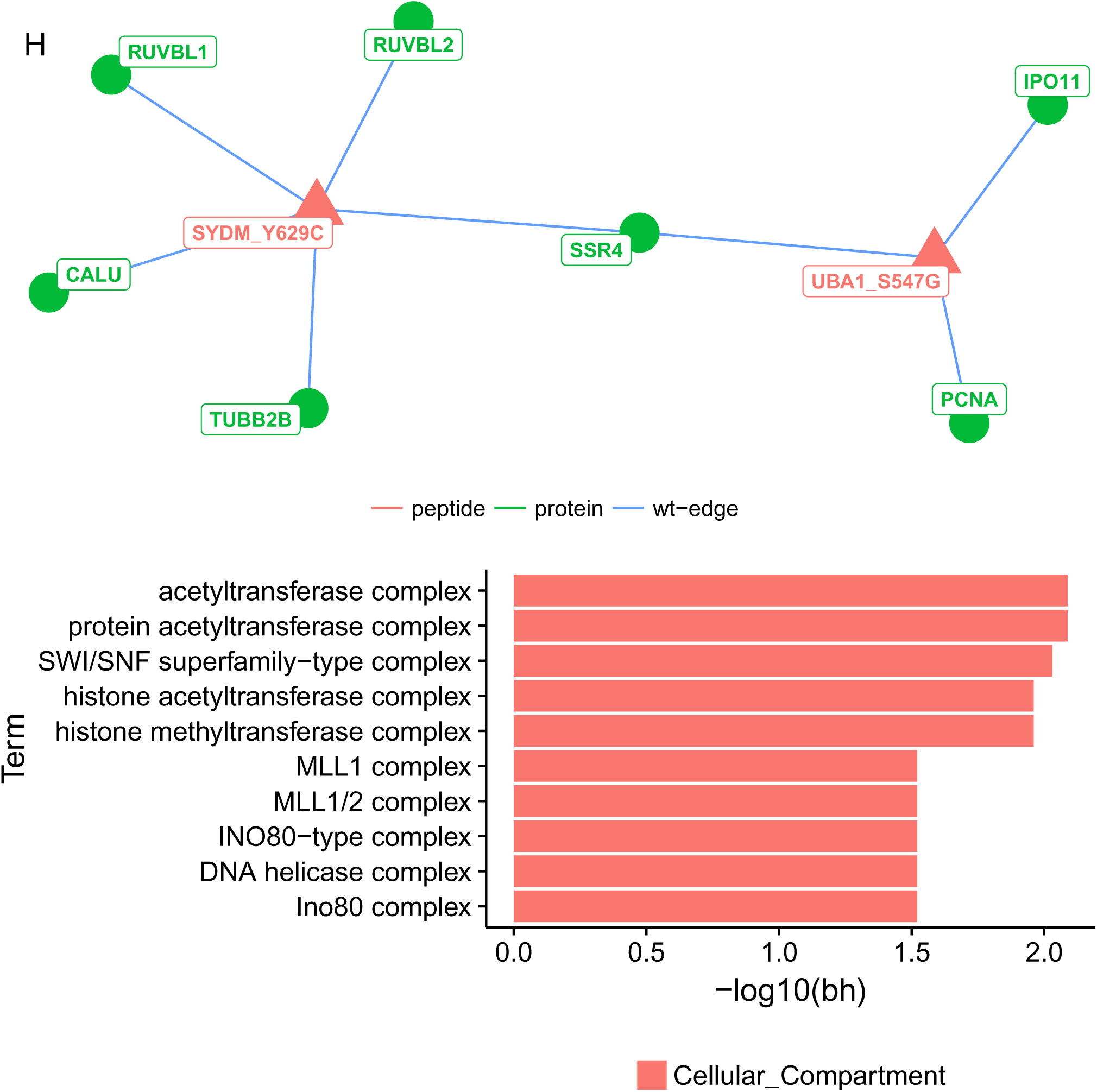
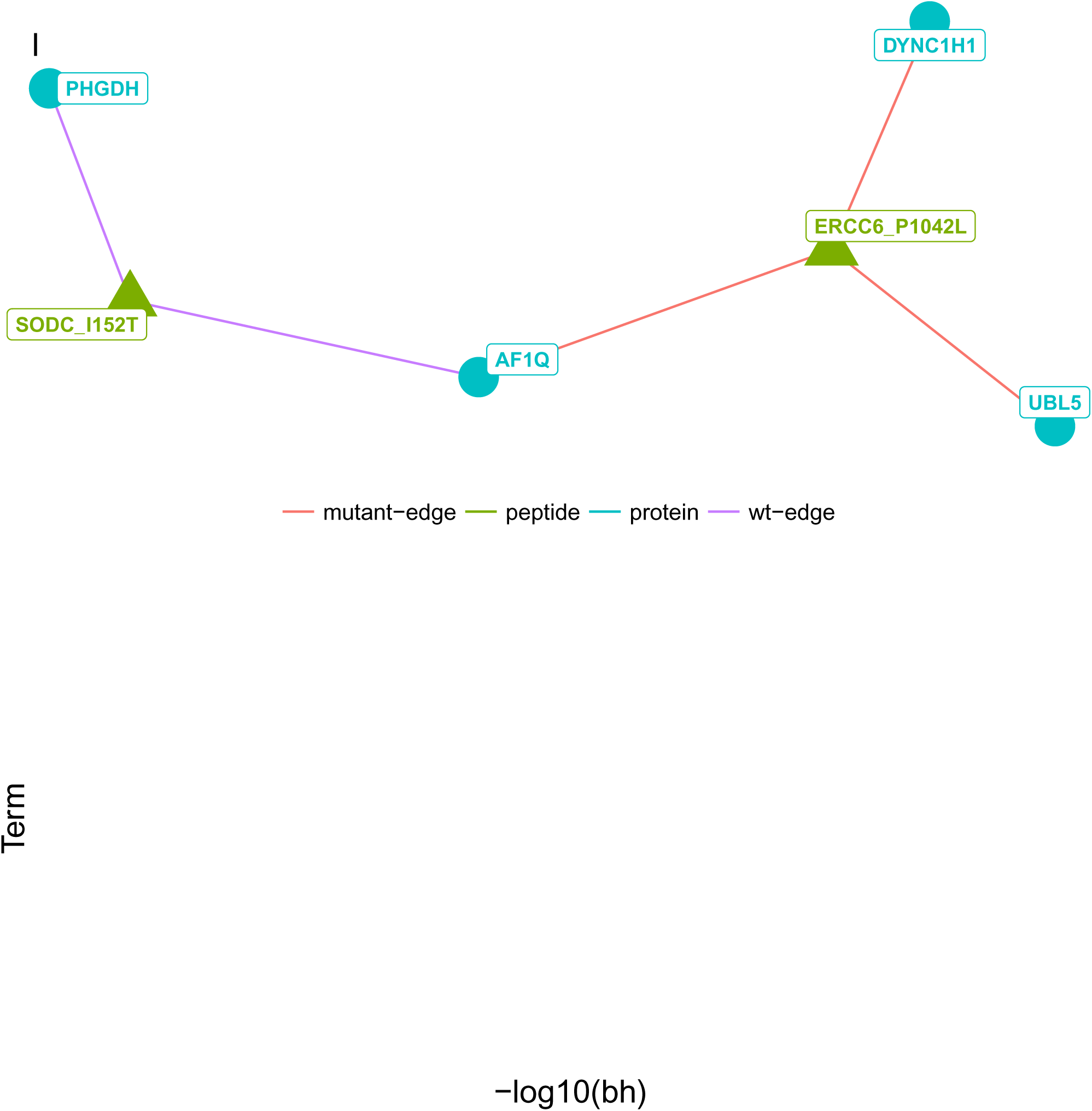
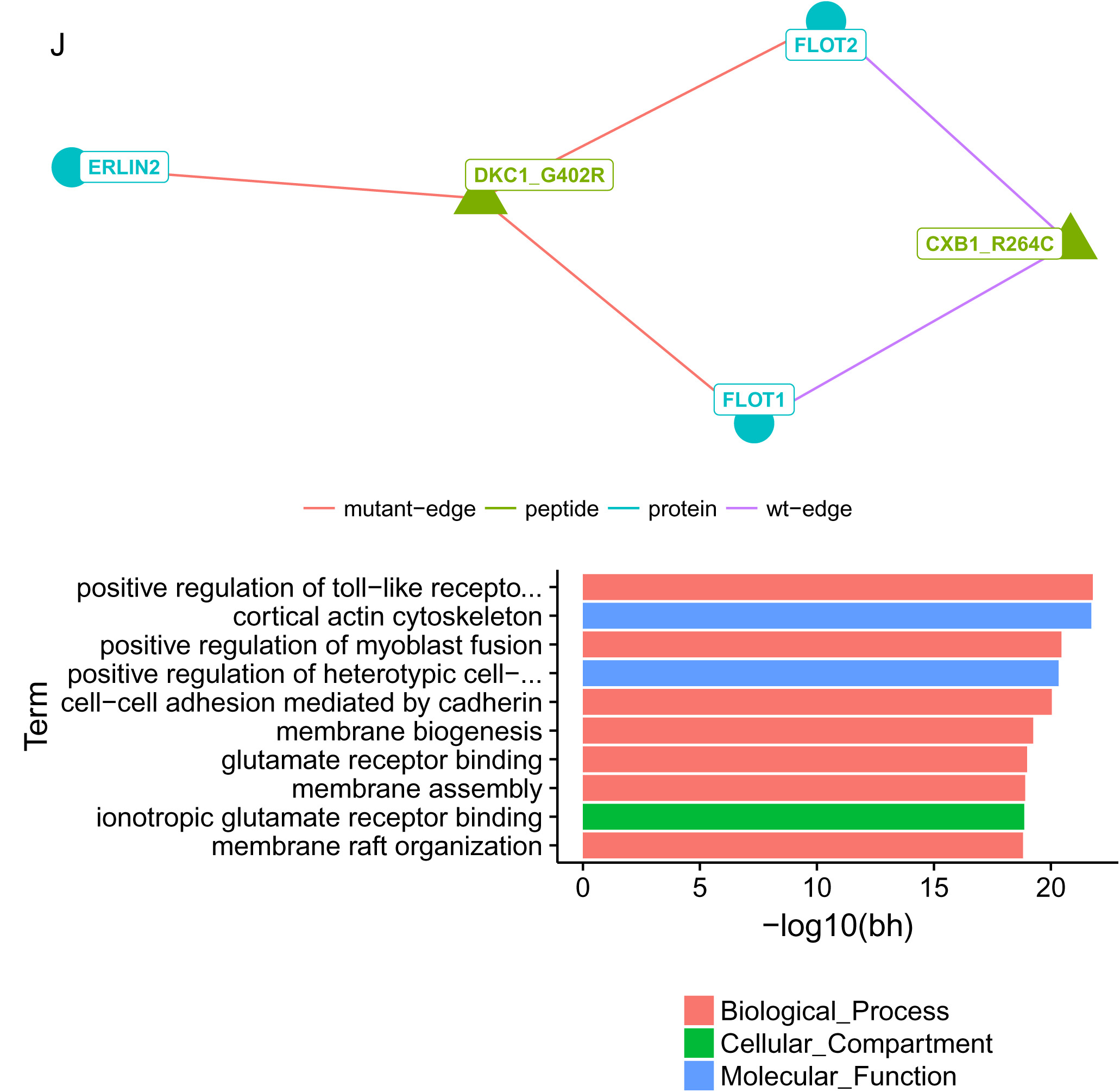
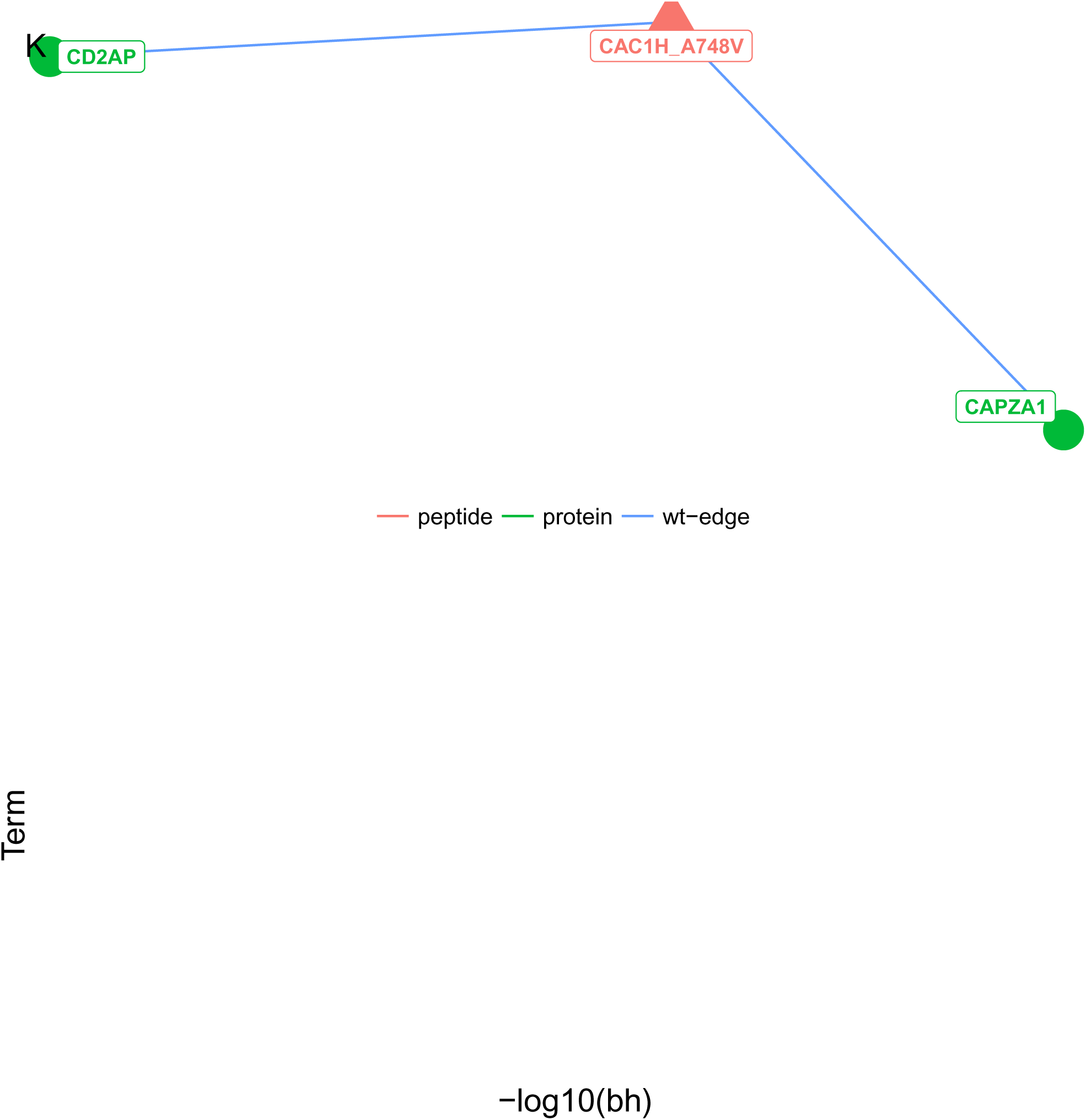
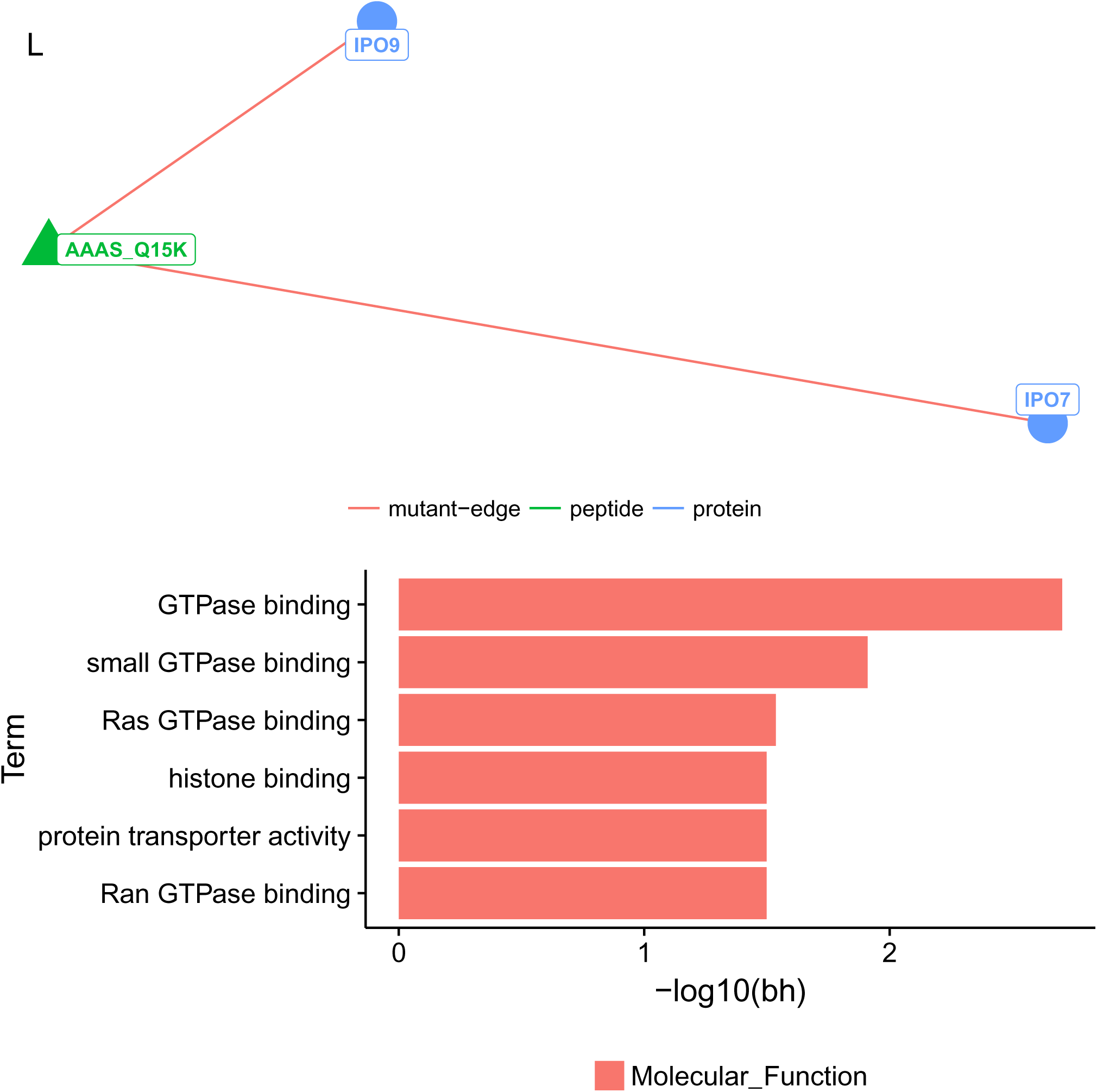
GO Term analysis of subclusters of the peptide-protein interaction network composed of significant differential interactions. 180 peptide-protein interactions that passed the strict LFQ filter and showed significant differential SILAC ratios between wild-type and mutant forms of the peptides were used to compose a peptide-protein interaction network. Communities (sub-graphs) of this network was extracted using fastgreedy.community function of the igraph R package (See Methods). GO term enrichment was calculated for the nodes of each subgraph of the network. Within the subgraphs, the edges (interactions) are colored differentially depending on whether the interaction is with a mutant or wild-type peptide. The peptides are depicted as triangles while the proteins are depicted as circles and they are differentially colored. Below the subgraph figure, a bar plot of the top 10 enriched GO terms are displayed (differentially colored by the GO term category as ‘Biological Process’, ‘Molecular Function’ or ‘Cellular Compartment), in which the x axis shows the log10 p-values corrected for multiple-testing correction using the Benjamini Hochberg method.

## References

1. Cooper, G. M. & Shendure, J. Needles in stacks of needles: finding disease-causal variants in a wealth of genomic data. Nat. Rev. Genet. 12, 628–640 (2011).

2. Subramanian, S. & Kumar, S. Evolutionary anatomies of positions and types of disease-associated and neutral amino acid mutations in the human genome. BMC Genomics 7, 306 (2006).

3. Yue, P., Li, Z. & Moult, J. Loss of protein structure stability as a major causative factor in monogenic disease. J. Mol. Biol. 353, 459–473 (2005).

4. Vacic, V. et al. Disease-associated mutations disrupt functionally important regions of intrinsic protein disorder. PLoS Comput. Biol. 8, e1002709 (2012).

5. Wright, P. E. & Jane Dyson, H. Intrinsically disordered proteins in cellular signalling and regulation. Nat. Rev. Mol. Cell Biol. 16, 18–29 (2014).

6. Uversky, V. N., Oldfield, C. J. & Dunker, A. K. Intrinsically disordered proteins in human diseases: introducing the D2 concept. Annu. Rev. Biophys. 37, 215–246 (2008).

7. Ryan, C. J. et al. High-resolution network biology: connecting sequence with function. Nat. Rev. Genet. 14, 865–879 (2013).

8. Wang, P. I. & Marcotte, E. M. It’s the machine that matters: Predicting gene function and phenotype from protein networks. J. Proteomics 73, 2277–2289 (2010).

9. Wang, X. et al. Three-dimensional reconstruction of protein networks provides insight into human genetic disease. Nat. Biotechnol. 30, 159–164 (2012).

10. Zhong, Q. et al. Edgetic perturbation models of human inherited disorders. Mol. Syst. Biol. 5, 321 (2009).

11. Hosp, F. et al. Quantitative interaction proteomics of neurodegenerative disease proteins. Cell Rep. 11, 1134–1146 (2015).

12. Van Roey, K. et al. Short linear motifs: ubiquitous and functionally diverse protein interaction modules directing cell regulation. Chem. Rev. 114, 6733–6778 (2014).

13. Fuxreiter, M., Tompa, P. & Simon, I. Local structural disorder imparts plasticity on linear motifs. Bioinformatics 23, 950–956 (2007).

14. Tompa, P., Davey, N. E., Gibson, T. J. & Babu, M. M. A million peptide motifs for the molecular biologist. Mol. Cell 55, 161–169 (2014).

15. Kadaveru, K., Vyas, J. & Schiller, M. R. Viral infection and human disease--insights from minimotifs. Front. Biosci. 13, 6455–6471 (2008).

16. Silvis, M. R. et al. A mutation in the cystic fibrosis transmembrane conductance regulator generates a novel internalization sequence and enhances endocytic rates. J. Biol. Chem. 278, 11554–11560 (2003).

17. Cordeddu, V. et al. Mutation of SHOC2 promotes aberrant protein N-myristoylation and causes Noonan-like syndrome with loose anagen hair. Nat. Genet. 41, 1022–1026 (2009).

18. Vogt, G. et al. Gains of glycosylation comprise an unexpectedly large group of pathogenic mutations. Nat. Genet. 37, 692–700 (2005).

19. Radivojac, P. et al. Gain and loss of phosphorylation sites in human cancer. Bioinformatics 24, i241–i247 (2008).

20. Narayan, S., Bader, G. D. & Reimand, J. Frequent mutations in acetylation and ubiquitination sites suggest novel driver mechanisms of cancer. Genome Med. 8, 55 (2016).

21. Uyar, B., Weatheritt, R. J., Dinkel, H., Davey, N. E. & Gibson, T. J. Proteome-wide analysis of human disease mutations in short linear motifs: neglected players in cancer? Mol. Biosyst. 10, 2626–2642 (2014).

22. Neduva, V. & Russell, R. B. Linear motifs: Evolutionary interaction switches. FEBS Lett. 579, 3342–3345 (2005).

23. Schulze, W. X. & Mann, M. A novel proteomic screen for peptide-protein interactions. J. Biol. Chem. 279, 10756–10764 (2004).

24. Gingras, A.-C. & Raught, B. Beyond hairballs: The use of quantitative mass spectrometry data to understand protein-protein interactions. FEBS Lett. 586, 2723–2731 (2012).

25. Gstaiger, M. & Aebersold, R. Applying mass spectrometry-based proteomics to genetics, genomics and network biology. Nat. Rev. Genet. 10, 617–627 (2009).

26. Meyer, K. & Selbach, M. Quantitative affinity purification mass spectrometry: a versatile technology to study protein–protein interactions. Front. Genet. 6, (2015).

27. Smits, A. H. & Vermeulen, M. Characterizing Protein-Protein Interactions Using Mass Spectrometry: Challenges and Opportunities. Trends Biotechnol. 34, 825–834 (2016).

28. Cox, J. et al. Accurate proteome-wide label-free quantification by delayed normalization and maximal peptide ratio extraction, termed MaxLFQ. Mol. Cell. Proteomics 13, 2513–2526 (2014).

29. Mann, M. Functional and quantitative proteomics using SILAC. Nat. Rev. Mol. Cell Biol. 7, 952–958 (2006).

30. Deng, H., Gao, K. & Jankovic, J. The role of FUS gene variants in neurodegenerative diseases. Nat. Rev. Neurol. 10, 337–348 (2014).

31. Dormann, D. et al. ALS-associated fused in sarcoma (FUS) mutations disrupt Transportin-mediated nuclear import. EMBO J. 29, 2841–2857 (2010).

32. Rogelj, B. et al. Widespread binding of FUS along nascent RNA regulates alternative splicing in the brain. Sci. Rep. 2, 603 (2012).

33. Ishigaki, S. et al. Position-dependent FUS-RNA interactions regulate alternative splicing events and transcriptions. Sci. Rep. 2, 529 (2012).

34. Qiu, H. et al. ALS-associated mutation FUS-R521C causes DNA damage and RNA splicing defects. J. Clin. Invest. 124, 981–999 (2014).

35. Yang, L., Embree, L. J., Tsai, S. & Hickstein, D. D. Oncoprotein TLS Interacts with Serine-Arginine Proteins Involved in RNA Splicing. J. Biol. Chem. 273, 27761–27764 (1998).

36. McMahon, H. T. & Boucrot, E. Molecular mechanism and physiological functions of clathrin-mediated endocytosis. Nat. Rev. Mol. Cell Biol. 12, 517–533 (2011).

37. Ferguson, S. M. & De Camilli, P. Dynamin, a membrane-remodelling GTPase. Nat. Rev. Mol. Cell Biol. 13, 75–88 (2012).

38. Pandey, K. N. Functional roles of short sequence motifs in the endocytosis of membrane receptors. Front. Biosci. 14, 5339–5360 (2009).

39. Dinkel, H. et al. ELM 2016—data update and new functionality of the eukaryotic linear motif resource. Nucleic Acids Res. 44, D294–D300 (2015).

40. Staudt, C., Puissant, E. & Boonen, M. Subcellular Trafficking of Mammalian Lysosomal Proteins: An Extended View. Int. J. Mol. Sci. 18, (2016).

41. Traub, L. M. Tickets to ride: selecting cargo for clathrin-regulated internalization. Nat. Rev. Mol. Cell Biol. 10, 583–596 (2009).

42. De Vivo, D. C. et al. Defective Glucose Transport across the Blood-Brain Barrier as a Cause of Persistent Hypoglycorrhachia, Seizures, and Developmental Delay. N. Engl. J. Med. 325, 703–709 (1991).

43. Pascual, J. M. et al. Structural signatures and membrane helix 4 in GLUT1: inferences from human blood-brain glucose transport mutants. J. Biol. Chem. 283, 16732–16742 (2008).

44. Roux, K. J., Kim, D. I., Raida, M. & Burke, B. A promiscuous biotin ligase fusion protein identifies proximal and interacting proteins in mammalian cells. J. Cell Biol. 196, 801–810 (2012).

45. Gibson, T. J., Dinkel, H., Van Roey, K. & Diella, F. Experimental detection of short regulatory motifs in eukaryotic proteins: tips for good practice as well as for bad. Cell Commun. Signal. 13, (2015).

46. Raiborg, C. & Stenmark, H. The ESCRT machinery in endosomal sorting of ubiquitylated membrane proteins. Nature 458, 445–452 (2009).

## References

1. UniProt Consortium. Reorganizing the protein space at the Universal Protein Resource (UniProt). Nucleic Acids Res. 40, D71–5 (2012).

2. Famiglietti, M. L. et al. Genetic variations and diseases in UniProtKB/Swiss-Prot: the ins and outs of expert manual curation. Hum. Mutat. 35, 927–935(2014).

3. Dosztányi, Z., Csizmok, V., Tompa, P. & Simon, I. IUPred: web server for the prediction of intrinsically unstructured regions of proteins based on estimated energy content. Bioinformatics 21, 3433–3434(2005).

4. Köhler, S. et al. The Human Phenotype Ontology in 2017. Nucleic Acids Res. 45, D865–D876(2017).

5. Schwanhöusser, B. et al. Global quantification of mammalian gene expression control. Nature 473, 337–342(2011).

6. Rappsilber, J., Ishihama, Y. & Mann, M. Stop and go extraction tips for matrix-assisted laser desorption/ionization, nanoelectrospray, and LC/MS sample pretreatment in proteomics. Anal. Chem. 75, 663–670(2003).

7. Cox, J. & Mann, M. MaxQuant enables high peptide identification rates, individualized p.p.b.-range mass accuracies and proteome-wide protein quantification. Nat. Biotechnol. 26, 1367–1372(2008).

8. Keilhauer, E. C., Hein, M. Y. & Mann, M. Accurate protein complex retrieval by affinity enrichment mass spectrometry (AE-MS) rather than affinity purification mass spectrometry (AP-MS). Mol. Cell. Proteomics 14, 120–135(2015).

9. Schulze, W. X. & Mann, M. A Novel Proteomic Screen for Peptide-Protein Interactions. J. Biol. Chem. 279, 10756–10764(2003).

10. Slaughter, L., Vartzelis, G. & Arthur, T. New GLUT-1 mutation in a child with treatment-resistant epilepsy. Epilepsy Res. 84, 254–256(2009).

11. Couzens, A. L. et al. Protein interaction network of the mammalian Hippo pathway reveals mechanisms of kinase-phosphatase interactions. Sci. Signal. 6, rs15 (2013).

12. Schindelin, J. et al. Fiji: an open-source platform for biological-image analysis. Nat. Methods 9, 676–682(2012).

13. Dinkel, H. et al. ELM 2016--data update and new functionality of the eukaryotic linear motif resource. Nucleic Acids Res. 44, D294–300 (2016).

14. Finn, R. D. et al. The Pfam protein families database: towards a more sustainable future. Nucleic Acids Res. 44, D279–85 (2016).

15. Shannon, P. et al. Cytoscape: a software environment for integrated models of biomolecular interaction networks. Genome Res. 13, 2498–2504(2003).

16. Csardi, G. & Nepusz, T. The igraph software package for complex network research. InterJournal, Complex Systems 1695 (2006).

17. Briatte, F. ggnetwork: Geometries to Plot Networks with 'ggplot2', R package version 0.5.1. CRAN (2016).

18. Wilkinson, L. ggplot2: Elegant Graphics for Data Analysis by WICKHAM, H. Biometrics 67, 678–679(2011).

19. Alexa, A. & Rahnenfuhrer, J. topGO: Enrichment Analysis for Gene Ontology. R package version 2.24.0. CRAN (2016).

